# Connectome-constrained modeling identifies neurons and synapses that sustain spontaneous activity in *Drosophila*

**DOI:** 10.64898/2026.08.21.745055

**Authors:** Qianzhu Li, Wenhao Ping, Ke Zhang, Chaoming Wang

## Abstract

Synapse-resolution connectomes specify a brain’s wiring; brain-wide recordings capture its activity. Neither alone identifies the cells and synapses that generate the activity. We bridge them by fitting a FlyWire connectome-constrained whole-brain model to calcium recordings of spontaneous activity in head-fixed *Drosophila*, then probing it *in silico* at cellular and synaptic resolution. The fitted model reproduces three features it was never trained on: lognormal synaptic weights, scale-free neuronal avalanches, and short intrinsic time constants in visual cells, each consistent with experiment. Systematic perturbations of the fitted whole-brain model show that spontaneous activity is not distributed uniformly across the connectome, but is organized by a compact neuropil core. Within this core, a highly sparse, brain-spanning ensemble of inhibitory hub neurons and their reciprocal synapses with excitatory partners are necessary and sufficient to sustain whole-brain resting-state dynamics. Connectome-constrained modeling therefore converts wiring diagrams and recordings into a perturbable digital platform that identifies the cells and synapses sustaining resting-state dynamics.

## Introduction

How the wiring of a brain produces its time-varying activity is a central question in neuroscience. Whole-brain connectomes now specify, neuron by neuron and synapse by synapse, how an animal’s brain is wired^[1–3]^. Brain-wide calcium imaging and dense electrophysiology capture the activity that wiring produces^[4–6]^. Neither dataset alone identifies the cells and synapses responsible. A connectome lists connections but not the *in vivo* strengths at which they operate, nor how they combine to generate population dynamics; a recording shows what the brain does without naming the circuit elements that produced it. Joining the two requires a generative model that respects every connection in the wiring diagram and reproduces the activity it carries.

The adult *Drosophila* brain is the only system in which both datasets currently exist at matching resolution. FlyWire reconstructs the entire central brain at synaptic resolution, comprising ∼140,000 neurons and ∼15 million synaptic connections, with cell-type and neurotransmitter labels for every cell^[1, 7]^. Whole-brain calcium imaging in head-fixed flies records activity from the same volume during rest and behavior^[8, 9]^. Resting-state activity in the fly carries the macroscopic signatures observed across species: spatiotemporal coherence^[10, 11]^ and near-critical avalanche statistics^[12–14]^. Yet their cellular and synaptic substrate has never been identified in any species. The fly is therefore where a synapse-resolution mechanistic account of resting-state dynamics can be built and tested.

Here we fit a whole-brain dynamical model, constrained by the FlyWire synapse-resolution connectome^[1]^, to calcium recordings of spontaneous activity in head-fixed *Drosophila*^[9]^. We then interrogate the fitted model *in silico* at neuropil, cellular, and synaptic resolution. Fitting turns the connectome from a static wiring diagram into an anatomical hypothesis space in which circuit mechanisms can be tested by direct perturbation. Within it we can silence any neuron or synapse and read out the brain-wide consequence, including manipulations no present experiment can deliver. The fit uses spontaneous calcium dynamics alone, yet the model recovers three features it never saw during training: lognormal synaptic weights, scale-free neuronal avalanches, and short intrinsic time constants in visual neurons. Agreement with these independent measurements grounds the model biologically and supports its use as a digital perturbation platform.

Using this platform, we find that resting-state activity is not a uniform property of the connectome; a compact neuropil core organizes it. Within this core, the dynamics depend on a highly sparse, brain-spanning ensemble of inhibitory hub neurons, whose reciprocal synapses with excitatory partners are necessary and sufficient for sustaining resting-state activity. Silencing one inhibitory hub releases its excitatory partner into runaway activity, whereas co-silencing that partner restores the resting regime. The hub comprises a small, experimentally accessible set of cell types, including MBON06, LPi13, LPi15, etc. This whole-brain modeling thus opens a route from wiring diagrams and recordings to testable cellular and synaptic mechanisms of resting-state activity.

## Results

### Connectome-constrained resting-state fitting yields lognormal synaptic weights and scale-free avalanche dynamics

We built the model from the FlyWire v783 connectome (Fig. 1b)^[1]^. The wiring and each presynaptic neuron’s excitatory or inhibitory sign stayed fixed throughout. Every neuron followed a first-order rate equation driven by its connectome-defined inputs and by an independent noise source (Fig. 1c), and a fixed anatomical readout pooled the single-neuron rates into the 73 neuropil signals the recordings provide (Fig. 1d). Training adjusted four things: the strength of each synapse, each neuron’s time constant, and a global noise scale and offset. We fit them by gradient descent to firing rates deconvolved from whole-brain calcium recordings of head-fixed flies (Fig. 1a,e)^[8, 9]^, using the first half of each ∼14-minute trace for training and holding out the second. The loss fell monotonically to a stable plateau (Fig. 1f). Every result below comes from five independently trained models (*n* = 5 throughout), so *n* counts model replicates rather than flies.

**Figure 1.**
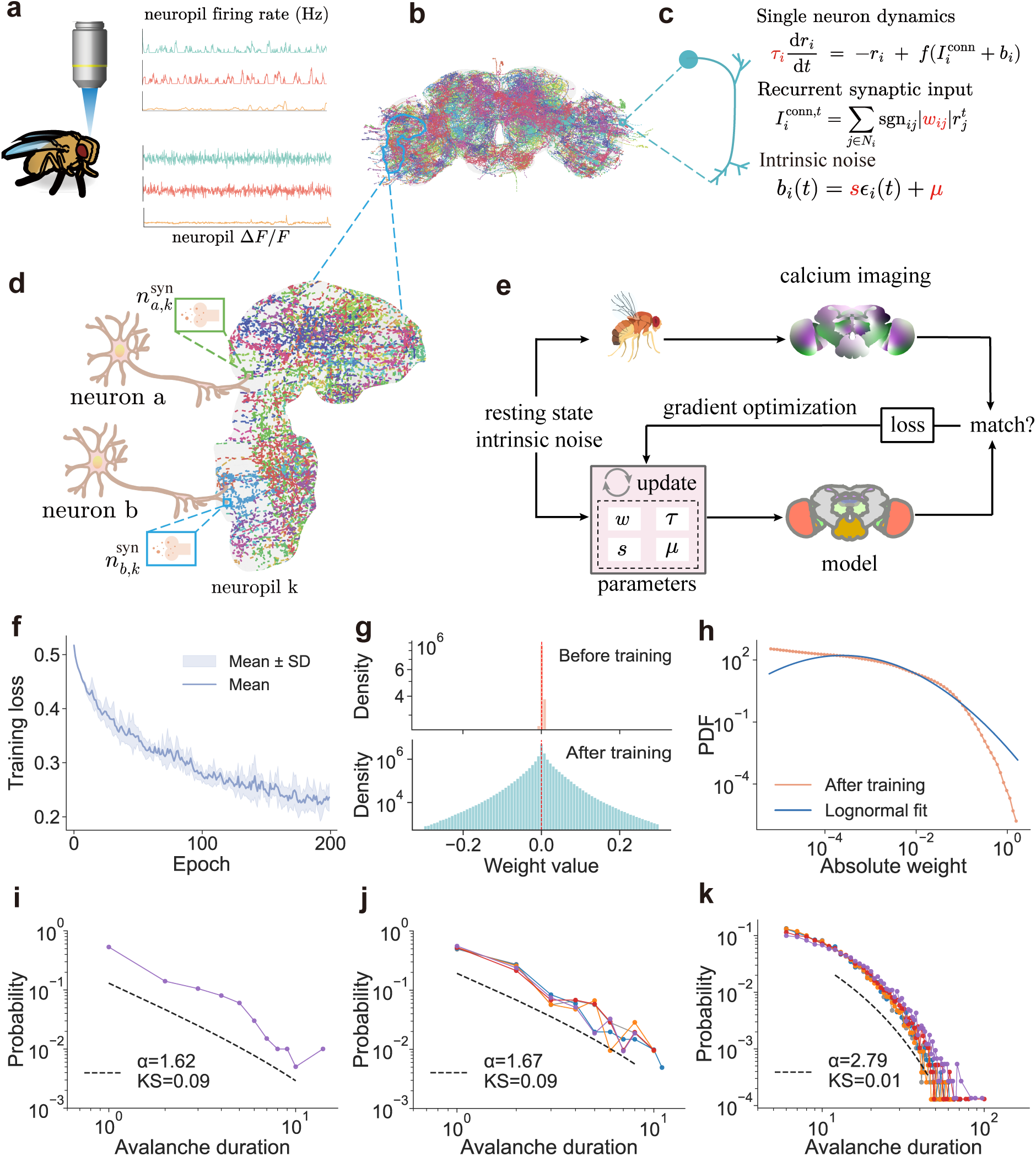
A connectome-constrained whole-brain model fits *Drosophila* resting-state dynamics and reproduces critical avalanche statistics. **a**, Whole-brain resting-state calcium imaging from head-fixed flies^[8, 9]^: neuropil Δ*F/F* (bottom) was deconvolved into population firing-rate estimates (top), which served as training targets. **b,** Structural scaffold derived from the FlyWire (v783) connectome (138,639 neurons; 15,091,983 chemical synapses); connection topology and synaptic polarity are fixed. **c,** Schematic of the single-neuron model. Each neuron integrates connectome-defined recurrent input from presynaptic partners *j* ∈ *N_i_* and an intrinsic noise drive *b_i_*(*t*) through a first-order rate equation with rectified-linear activation (full equation in Methods, Eq. (3)). **d,** Schematic of the connectome-derived neuropil readout: simulated single-neuron rates are aggregated into a population signal weighted by per-neuron synapse counts *n^syn^_i,k_* within each neuropil *k* (Methods, Eq. (6)). **e,** Calibration loop: free parameters (synaptic-weight magnitudes {|*w_ij_*|}, per-neuron time constants {*τ_i_*}, global noise scale *s*, and offset *µ*) are optimized by gradient descent to minimize the mean-squared error between simulated and empirical neuropil activity (Methods, Eq. (7)). **f,** Training loss over optimization epochs. Line and shaded band, mean ± std.; *n* = 5 independently trained models. **g,** Effective synaptic-weight distributions before (top) and after (bottom) training; *n* = 5 trained models pooled. **h,** Distribution of absolute synaptic weights after training (log–log axes; *n* = 5 trained models pooled, ∼ 1.5 × 10^7^ synapses each). Maximum-likelihood truncated lognormal fit: *µ* = −4.97, *σ* = 1.83, Kolmogorov–Smirnov *KS* = 0.06. Lognormal preferred over both power-law and exponential alternatives by Vuong-corrected likelihood-ratio tests; both *p* values fall below the double-precision floor (*p <* 10*^−^*^300^) at *N* ≈ 1.5 × 10^7^ synapses, and the underlying log-likelihood-ratio statistics are reported in Supplementary Note 3. **i,** Avalanche-duration distribution from the empirical neuropil-level calcium imaging trace (single example fly). Maximum-likelihood truncated power-law fit: exponent *α* = 1.62, *KS* = 0.09; truncated power-law preferred over the lognormal alternative by a Vuong-corrected likelihood-ratio test (*p* = 2.26 × 10*^−^*^7^); the exponential alternative is rejected with smaller *p* (Supplementary Note 3). **j,** Avalanche-duration distribution from simulated neuropil activity (*n* = 5 independently trained models, events pooled for the fit). Maximum-likelihood truncated power-law fit: *α* = 1.67, *KS* = 0.09; truncated power-law preferred over the lognormal alternative by a Vuong-corrected likelihood-ratio test (*p* = 6.31 × 10*^−^*^3^); the exponential alternative is rejected with smaller *p* (Supplementary Note 3). **k,** Single-neuron avalanche-duration distribution from simulated activity (*n* = 5 independently trained models, events pooled for the fit). Maximum-likelihood truncated power-law fit: *α* = 2.79, *KS* = 0.01; truncated power-law preferred over the lognormal alternative by a Vuong-corrected likelihood-ratio test (*p* = 2.98 × 10*^−^*^10^); the exponential alternative is rejected with smaller *p* (Supplementary Note 3). Neuropil abbreviations follow the Ito reference atlas^[15]^; see Methods.

We then asked what the fitted model produced beyond its training target. The synaptic weights gave the first test. Initialization placed every magnitude in a narrow peak; training spread them over several orders of magnitude on both the excitatory and the inhibitory side (Fig. 1g, Supplementary Fig. 1). The fitted distribution is lognormal (Fig. 1h), with power-law and exponential alternatives rejected. Lognormal weights recur across cortical microcircuits^[16]^. Here one emerged from fitting activity alone; nothing in the loss asked for it.

Neuronal avalanches gave a second test. Spontaneous activity across species follows power-law distributions of avalanche size and duration^[12–14, 17]^. The recordings carried this signature and the simulated neuropil activity reproduced it, with matching exponents (durations, *α_D_* = 1.62 empirical versus 1.67 simulated; sizes, *α_S_* = 1.30 versus 1.38; Fig. 1i,j, Supplementary Fig. 2a,b)^[8, 9]^. Both satisfy the crackling-noise scaling relation (*α_S_* − 1)*/*(*α_D_* − 1), at 0.48 and 0.57, inside the range reported for near-critical cortical recordings^[18, 19]^. The model also lets us count avalanches one neuron at a time, which no current recording allows. Exponents there were steeper (Fig. 1k, Supplementary Fig. 2c), as expected when an avalanche is a per-neuron burst rather than a network-wide cascade, and consistent with the steepening reported at fine spatial scales^[20, 21]^.

### Connectome structural priors are required for stable and generalizable resting-state dynamics

Does the connectome scaffold do this work, or would any recurrent network with enough parameters? We trained a gated recurrent unit (GRU) with a matched parameter count on the same loss (Supplementary Fig. 3). Both models fit the training half well: their functional connectivity (FC) matched the recordings at *r* = 0.994 for the connectome-constrained model and *r* = 0.941 for the GRU (Fig. 2c–e). The held-out half separated them. Resting-state FC drifts even within the recordings, so the empirical train–test consistency of *r* = 0.473 sets a ceiling on what any model can match. The connectome-constrained model reached *r* = 0.423 against the test-set recordings, close to that ceiling; the GRU reached *r* = 0.288 (Fig. 2f–h). The simulated traces show the same split. The connectome-constrained model kept generating stable, irregular activity through the test segment (Fig. 2a), whereas the GRU lost the fast fluctuations of the recordings, some neuropils decaying toward quiescence and others drifting into slow, large excursions (Fig. 2b). Generalizing beyond the training window needs a structural prior, not more capacity.

**Figure 2.**
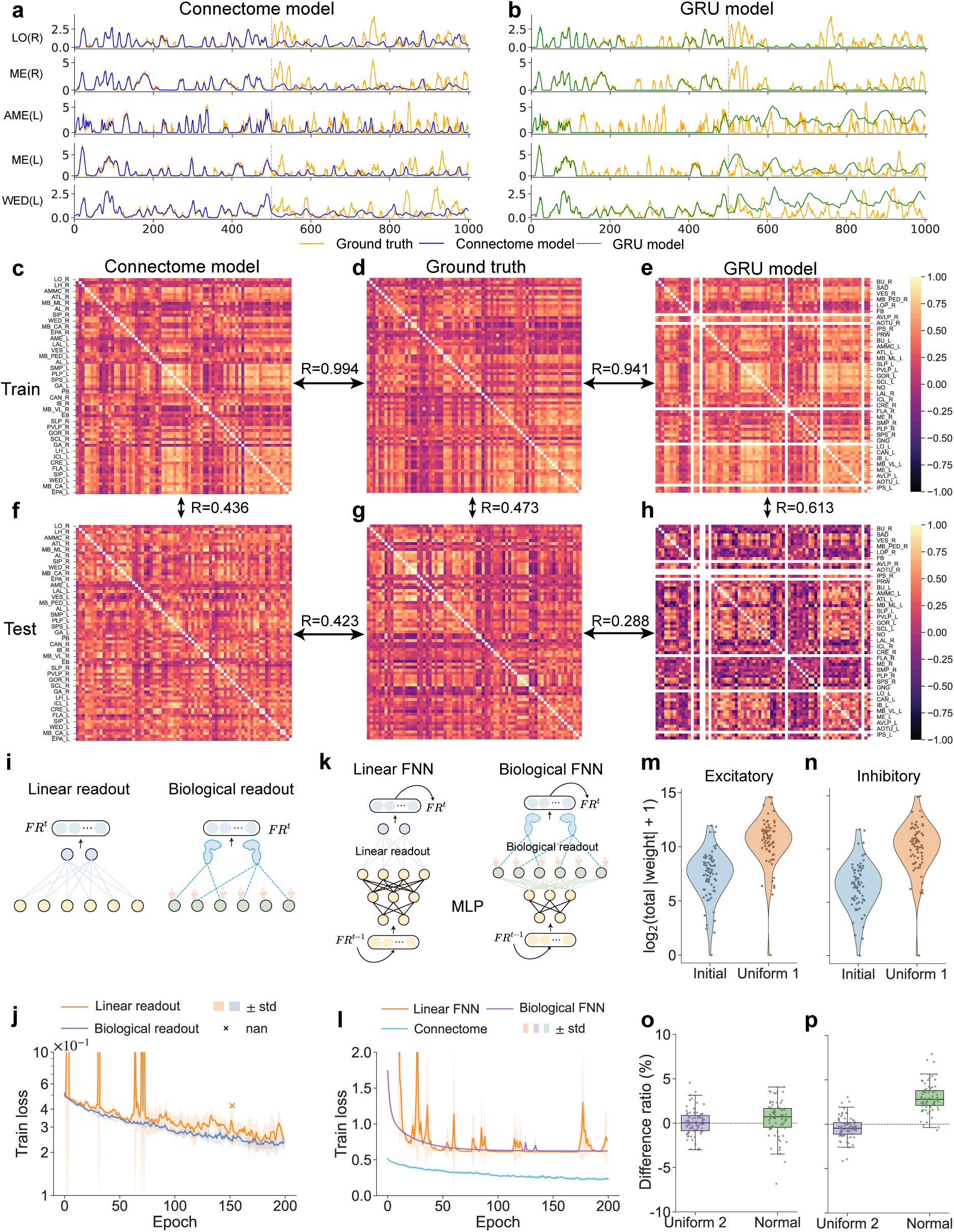
Biological connectome constraints are essential for accurate and stable whole-brain resting-state modeling. **a,b**, Simulated activity of five representative neuropils (LO(R), ME(R), AME(L), ME(L), WED(L)) after convergence, for the connectome-constrained model (**a**, blue) and a parameter-matched gated recurrent unit (GRU) model (**b**, green), each overlaid on empirical ground truth (orange). Vertical dashed line marks the boundary between the training set (left) and the held-out test set (right). **c–e,** Functional connectivity (FC) matrices computed on the training set for the connectome-constrained model (**c**), the empirical recordings (**d**), and the GRU model (**e**). Horizontal arrows annotate Pearson correlations between adjacent matrices (*r* = 0.994 for **c**–**d**; *r* = 0.941 for **d**–**e**). **f–h,** FC matrices computed on the held-out test set for the connectome-constrained model (**f**), the empirical recordings (**g**), and the GRU model (**h**). Vertical arrows annotate the train–test consistency of each FC matrix (connectome-constrained model, *r* = 0.436; empirical, *r* = 0.473; GRU, *r* = 0.613); horizontal arrows annotate the model–empirical agreement on the test set (connectome-constrained model, *r* = 0.423; GRU, *r* = 0.288). **i,** Schematic of the two readout architectures compared in the recurrent setting: an unconstrained linear readout (left) versus the connectome-derived synapse-density-weighted readout (right). **j,** Training-loss curves over epochs for the connectome-constrained model with the unconstrained linear readout (orange) versus the connectome-derived readout (blue). Line and shading, mean ± std.; *n* = 5 independently trained models per condition; crosses mark divergent (NaN) runs. **k,** Schematic of the two feedforward control architectures trained under the next-step prediction paradigm: a multilayer perceptron (MLP) with an unconstrained linear readout (left) versus an MLP with the connectome-derived readout (right). **l,** Training-loss curves for the two feedforward controls in **k** (MLP with linear readout, orange; MLP with connectome-derived readout, purple) alongside the connectome-constrained recurrent model (blue). Line and shading, mean ± std.; *n* = 5 independently trained models per condition. **m,n,** Distributions of log_10_(total post-synaptic weight) per neuropil for excitatory (**m**) and inhibitory (**n**) populations, comparing the initial values (“Initial”, blue) and the converged values after training under a uniform prior (“Uniform 1”, orange). Violins show kernel density estimates pooled across *n* = 5 trained models; dots are individual neuropils (*N* = 73). **o,p,** Percentage difference between converged weights obtained under a second uniform prior (“Uniform 2”) or a truncated-normal prior (“Normal”) and those obtained under Uniform 1, plotted per neuropil for excitatory (**o**) and inhibitory (**p**) populations. Box plots: center line, mean; box, mean ± std.; whiskers, range; *n* = 5 independently trained models per prior, *N* = 73 neuropils. Neuropil abbreviations follow the Ito reference atlas^[15]^; see Methods.

Two ablations isolate which parts of that prior matter. Replacing the connectome-derived readout with a freely learned linear projection destabilized training, several runs diverging before reaching a usable fit (Fig. 2i,j). The anatomical readout is load-bearing, not decorative. Recurrence is the second prior. We tested it with two feedforward controls trained to predict the next time step: a parameter-matched multilayer perceptron (MLP), and the same MLP with the fixed connectome readout appended. Both plateaued at a higher loss than the recurrent model (Fig. 2k,l). Resting-state activity does not reduce to a one-step transition map.

Could the fit instead be an artifact of where optimization started? We retrained from three initial weight priors: a uniform interval (“Uniform 1”), a second uniform interval with different random seeds (“Uniform 2”), and a truncated normal centered at zero (“Normal”). The converged neuropil-level excitatory and inhibitory weights moved well away from their initial values (Fig. 2m,n; Supplementary Fig. 4a), and the three priors converged to within ±5% of one another (Fig. 2o,p; Supplementary Fig. 4b,c). The fitted parameters are reproducible across initializations, not artifacts of any one prior (Supplementary Note 4).

### A compact 14-neuropil core is necessary and sufficient in the model to sustain whole-brain resting-state dynamics

We then treated the fitted model as a preparation to be perturbed, starting at the neuropil level. One neuropil at a time, we silenced the outgoing synapses of all its neurons and measured how the prediction loss changed at every other neuropil (Fig. 3a). The resulting directed 73 × 73 effectome matrix is far from uniform (Fig. 3b). A few source neuropils disrupt activity brain-wide, most affect only a handful of targets, and a separate set of targets responds to ablations originating almost anywhere. Resting-state coupling is not a sum of independent pairwise interactions: a small set of source neuropils sets the brain-wide dynamics, and the rest follow.

**Figure 3.**
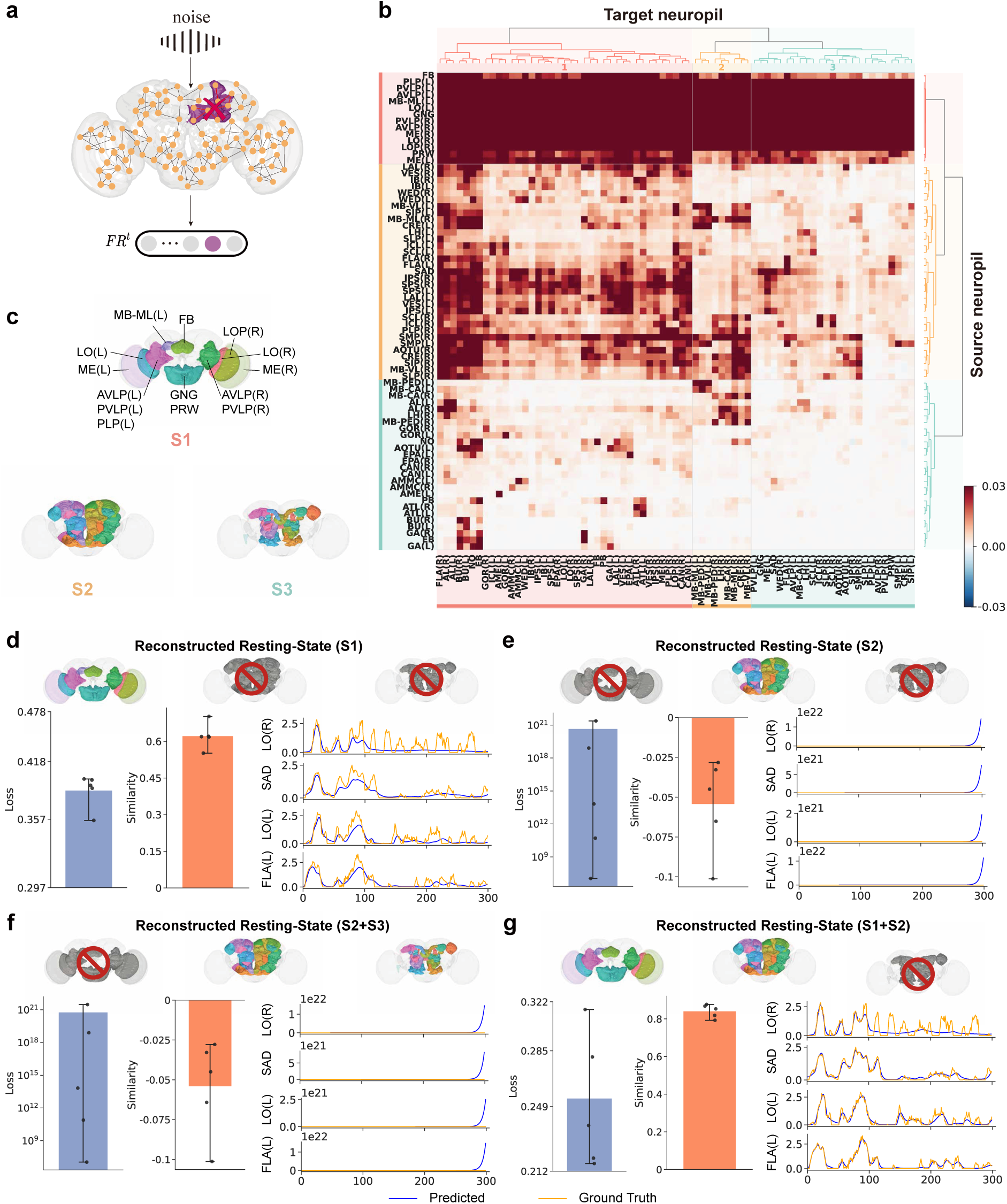
A neuropil-level effectome identifies a core subnetwork organizing whole-brain resting-state dynamics. **a**, Schematic of the neuropil-level effectome analysis: for each source neuropil, outgoing synapses of all constituent neurons were ablated *in silico* and the resulting change in prediction loss was measured at every target neuropil (Methods). **b,** Directed 73 × 73 effectome matrix of signed loss changes. Rows, source neuropils (ablation site); columns, target neuropils (readout); color, signed loss change (red, increase; blue, decrease; clipped at ±0.03). Colored side bars indicate source-cluster (right) and target-cluster (top) assignments from hierarchical agglomerative clustering (*k* = 3 for both, selected by silhouette criterion; Methods, Supplementary Fig. 5); accompanying dendrograms show the corresponding hierarchical relationships. The *k* = 3 partition is reproducible across the *n* = 5 trained models (Supplementary Fig. 6). **c,** Anatomical layout of the three principal source clusters identified by clustering the rows of **b**, projected onto the standard *Drosophila* brain atlas. Source cluster 1 (S1) is the most compact set of 14 neuropils, spanning visual-associated regions (ME(L), ME(R), LO(L), LO(R), LOP(R)), associative centers (FB, MB-ML(L)), and ventromedial neuropils (AVLP(L/R), PVLP(L/R), PLP(L), GNG, PRW); source cluster 2 (S2, 33 neuropils) and source cluster 3 (S3, 26 neuropils) span larger, more distributed neuropil groups. Full membership lists in Supplementary Table 1. **d–g,** Selective preservation of source clusters: in each condition, the outgoing synapses of the indicated cluster(s) are retained while the remainder are ablated *in silico*. Each panel reports test-set prediction loss summed over all neuropils (left, blue bar), Pearson correlation between simulated and empirical neuropil activity (middle, orange bar), and representative simulated traces for four example neuropils (LO(R), SAD, LO(L), FLA(L); predicted in color, empirical ground truth in gray). Brain icons indicate intact and ablated clusters (red prohibition symbol). **d,** S1 alone preserved. **e,** S2 alone preserved. **f,** S2+S3 preserved (S1 ablated). **g,** S1+S2 preserved (S3 ablated). With all clusters intact the model reaches loss ∼0.25 and *r* ∼ 0.8, the baseline referred to in the text; preserving S1+S3 instead of S1+S2 gives loss ∼0.38 and *r* ∼ 0.64 (Supplementary Fig. 8b). In **e** and **f** the simulated rates leave the bounded regime and the loss reaches 10^21^–10^22^; losses above 10^16^ indicate numerical divergence rather than a graded fit error, and their magnitudes are not comparable across conditions (Methods). Bars, mean ± std.; *n* = 5 independently trained models (one independent simulation per model per condition). Neuropil abbreviations follow the Ito reference atlas^[15]^; see Methods.

Clustering the rows of the matrix returned three source clusters, each a spatially compact anatomical set (S1, S2, S3; Fig. 3c). S1 is by far the smallest, 14 of the 73 neuropils, spanning visual-associated, central-complex, and inferior regions. S2 (33 neuropils) and S3 (26 neuropils) cover much more of the brain yet produce smaller brain-wide effects (full membership in Supplementary Table 1). A disproportionate share of the brain’s causal weight sits in the compact cluster.

Selective preservation tests then asked whether S1 is necessary and sufficient. We kept the outgoing synapses of one or more clusters, silenced the rest, and recorded prediction loss, Pearson correlation to the recordings, and simulated traces (Fig. 3d–g; whole-brain traces in Supplementary Fig. 7). With S1 alone preserved, the model still tracked the gross temporal envelope, at degraded accuracy: loss rose from ∼0.25 to ∼0.36 and *r* fell from ∼0.8 to ∼0.6 (Fig. 3d). Removing S1 was different in kind. Preserving only S2, only S3, or S2 and S3 together pushed firing rates out of the bounded regime entirely (Fig. 3e,f; Supplementary Fig. 8a). Adding S2 back to an intact S1 returned loss and correlation to baseline (Fig. 3g), and adding S3 instead worked nearly as well (Supplementary Fig. 8b), so S2 and S3 are interchangeable once S1 is present. S1 is necessary for bounded dynamics and sufficient for the coarse temporal structure of resting-state activity.

Clustering the columns gives the mirror image (Supplementary Fig. 9; Supplementary Table 1). Of the three target clusters, T1 (39 neuropils, optic, central-complex, and ventromedial) carries essentially all of the model’s simulated power, concentrated below ∼0.1 Hz, while T2 (9 neuropils, almost all mushroom-body subcompartments) and T3 (25 neuropils, superior and lateral protocerebrum) are spectrally flat. Ablating T1 *in silico* removes the low-frequency signature at every bin; ablating T2 or T3 leaves it intact.

S1 generates the slow signal and T1 expresses it: one low-rank source–target hub seen from two orthogonal axes. Mammalian connectomes contain the same kind of structure, a small ensemble of densely interconnected regions, the rich club, that disproportionately governs global brain states^[22–24]^. FlyWire network analysis finds it in *Drosophila* as well^[1, 25]^, and calcium imaging shows resting-state co-fluctuations organizing into reproducible modules centered on higher-order neuropils^[8, 9, 26]^. Individual members of S1 carry independent experimental support: LOP and lobula circuits sustain internally generated activity without sensory input^[27, 28]^; the dorsal fan-shaped body gates quiescence and sensory disengagement during sleep^[29, 30]^; and 20–30 Hz oscillations in the central brain, mushroom-body region included, track arousal and attentional state^[31]^.

### A small inhibitory hub population sustains resting-state dynamics in the model

Which neurons inside this core carry the dynamics? The model gives two handles on each cell: its time constant *τ* and its effective synaptic weights (Fig. 1c). Training left most of the 138,639 time constants near their starting values, but moved a minority sharply in both directions (Fig. 4a). We took the two tails: the 3% of neurons whose *τ* shortened most, and the 3% whose *τ* lengthened most. Projected onto the standard fly brain atlas^[15]^, both tails land in spatially compact, hemisphere-biased clusters rather than spreading uniformly (Supplementary Fig. 10). Timescale remodeling is anatomically selective.

**Figure 4.**
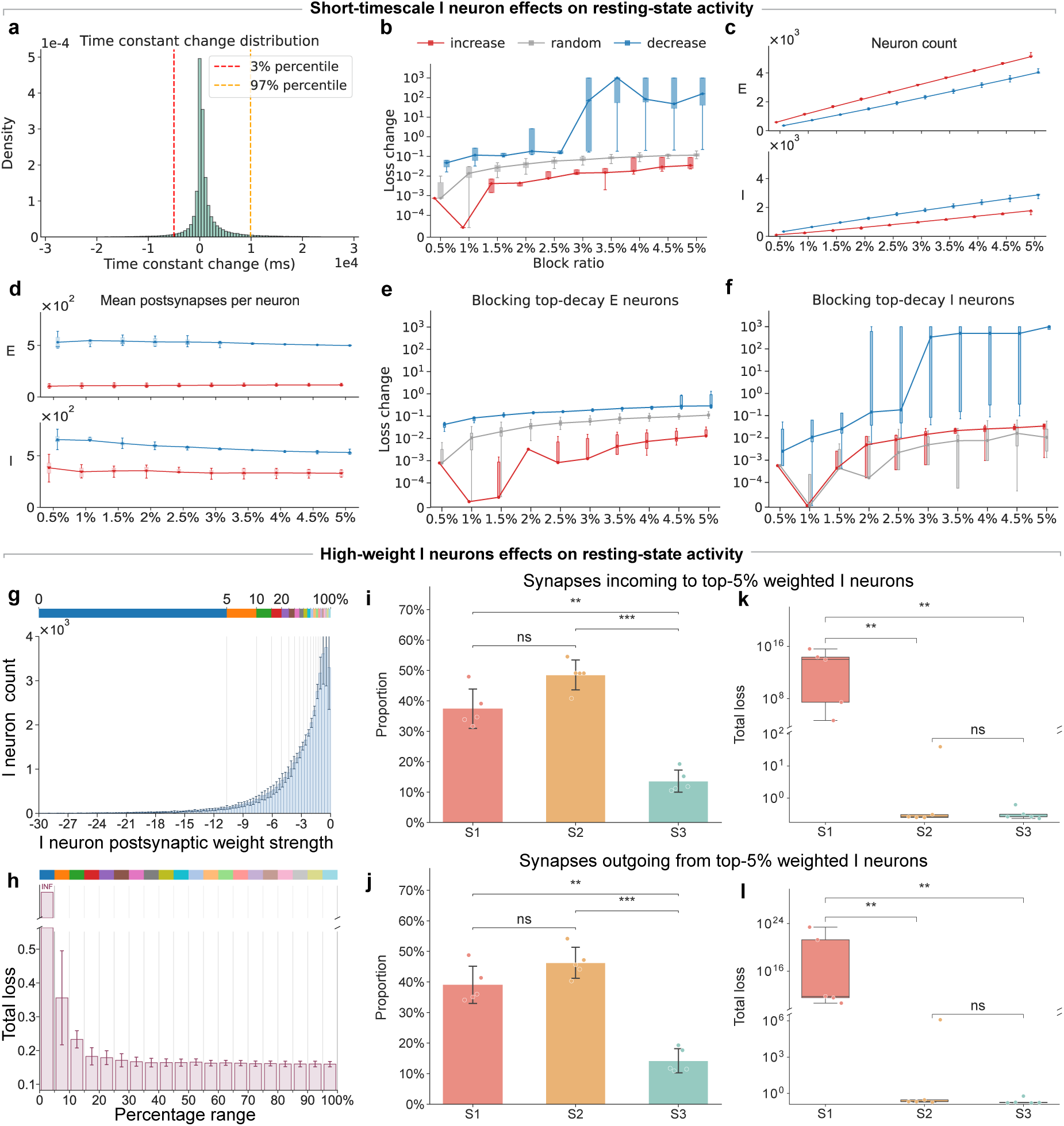
Essential short-timescale, high-weight inhibitory neurons in source cluster 1 sustain resting-state dynamics. **a**, Distribution of training-induced changes in the neuronal time constant (Δ*τ*) across all 138,639 neurons (mean across *n* = 5 trained models). Vertical dashed lines mark the 3rd and 97th percentile cutoffs used to define the top-3% negative tail (largest Δ*τ* decrease) and top-3% positive tail (largest Δ*τ* increase). **b,** Loss change resulting from *in silico* silencing of Δ*τ*-tail neurons across block ratios (0.5%–5%): Δ*τ*-decrease tail (blue), Δ*τ*-increase tail (red), matched random controls (gray, 10 draws per condition). For the decrease tail the loss stays near 10*^−^*^1^ up to a 2.5% block ratio and reaches ∼ 10^2^ at 3%. **c,** Cellular composition of the Δ*τ*-tail subsets across the 0.5%–5% block-ratio range: counts of excitatory (E, top) and inhibitory (I, bottom) neurons in the increase tail (red) and decrease tail (blue). **d,** Mean number of postsynapses per neuron for the Δ*τ*-tail excitatory (E, top) and inhibitory (I, bottom) subsets across the 0.5%–5% block-ratio range, shown separately for the increase tail (red) and decrease tail (blue). **e,** As in **b**, with silencing restricted to excitatory neurons. **f,** As in **b**, with silencing restricted to inhibitory neurons. Dashed blue, Δ*τ*-decrease inhibitory subset, for which the loss rises from ∼ 10*^−^*^1^ at a 2.5% block ratio to ∼ 10^3^ at 3%. **g,** Distribution of post-synaptic weight strength of inhibitory (I) neurons in the trained model, ranked by outgoing weight magnitude |*W^out^_i_*| (Methods, Eq. (11)). The colored ribbon at the top bins I neurons by percentile of post-synaptic weight magnitude (top 5%, 5–10%, 10–20%, …, 90–100%); *n* = 5 trained models pooled. **h,** Total loss when the I neurons in each percentile bin of **g** are silenced *in silico*; silencing the top-5% bin (largest post-synaptic weight magnitude) drives the loss to effectively unbounded values, dwarfing every other bin. Bars, mean; *n* = 5 trained models. **i,** Source-cluster composition of the synapses projecting to the top-5% weighted I neurons (clusters defined in Fig. 3c): ∼38% S1, ∼48% S2, ∼15% S3. Two-sided paired Student’s *t*-tests across *n* = 5 independently trained models with Holm–Bonferroni correction: S1 vs S2, *p* = 8.59 × 10*^−^*^2^, n.s.; S1 vs S3, *p* = 4.17 × 10*^−^*^3^; S2 vs S3, *p* = 1.98 × 10*^−^*^4^. **j,** Total loss (log_10_) when the incoming synapses to the top-5% weighted I neurons are blocked separately within each source cluster. Two-sided paired *t*-tests with Holm–Bonferroni correction: S1 vs S2, *p* = 6.79 × 10*^−^*^3^; S1 vs S3, *p* = 2.04 × 10*^−^*^3^; S2 vs S3, *p* = 0.383, n.s. **k,** Source-cluster composition of the synapses that the top-5% weighted I neurons project to: ∼40% S1, ∼45% S2, ∼15% S3. Two-sided paired *t*-tests with Holm–Bonferroni correction: S1 vs S2, *p* = 0.192, n.s.; S1 vs S3, *p* = 3.52 × 10*^−^*^3^; S2 vs S3, *p* = 4.86 × 10*^−^*^4^. **l,** Total loss (log_10_) when the outgoing synapses from the top-5% weighted I neurons are blocked separately within each source cluster. Two-sided paired *t*-tests with Holm–Bonferroni correction: S1 vs S2, *p* = 9.11 × 10*^−^*^3^; S1 vs S3, *p* = 5.34 × 10*^−^*^3^; S2 vs S3, *p* = 0.491, n.s. Box plots in **j,l**: center line, median; box, interquartile range (IQR); whiskers, 1.5×IQR; outliers shown as points; *n* = 5 independently trained models. Bars elsewhere, mean ± std. across *n* = 5 trained models. n.s. indicates *p* ≥ 0.05; exact corrected *p* values are reported above.

Silencing shows which tail matters. We silenced the shortened tail, the lengthened tail, or a matched random subset, sweeping the fraction of neurons removed from 0.5% to 5% (Fig. 4b). The shortened tail dominated at every fraction, by orders of magnitude. Its effect was also a threshold rather than a gradient: loss stayed near 10*^−^*^1^ up to a 2.5% block ratio, then jumped about three orders of magnitude at 3% and kept rising. Neurons whose intrinsic timescales training shortened are required for resting-state activity in the model; the lengthened-timescale neurons are largely dispensable.

What separates the two tails? The shortened tail holds more inhibitory neurons (Fig. 4c) and its neurons form more postsynapses (Fig. 4d). These are high-connectivity, inhibition-enriched cells, not generic ones. Repeating the sweep within each polarity locates the effect. Silencing excitatory neurons alone preserved the ordering (shortened *>* random *>* lengthened) but shrank every loss change by orders of magnitude (Fig. 4e). Silencing inhibitory neurons alone reproduced the full effect, threshold included (Fig. 4f). The shortened-timescale inhibitory neurons account for almost all of the disruption.

The Δ*τ* criterion depends on the optimization trajectory. To recover the same set from the trained weights alone, we ranked all inhibitory (I) neurons by outgoing weight magnitude |*W^out^_i_*| and silenced them one percentile bin at a time (Fig. 4g). Only the top-5% bin drove the loss to effectively unbounded values; every other bin sat orders of magnitude below it (Fig. 4h). Two independent criteria converge on one small inhibitory population.

Where do these neurons connect? Their incoming synapses are spread across all three source clusters, with S1 and S2 carrying statistically indistinguishable shares (Fig. 4i), and their outgoing synapses follow the same split (Fig. 4k). Blocking those synapses cluster by cluster is not symmetric. Blocking the S1 inputs destabilized the rate dynamics, while matched S2 or S3 inputs left the model near baseline (Fig. 4j); only the S1-routed outputs did the same on the output side (Fig. 4l). Anatomical share does not predict causal weight. The top-5% weighted inhibitory neurons and their reciprocal synapses routed through S1 form the compact synaptic substrate of the model’s resting-state dynamics.

### Reciprocal excitatory–inhibitory interactions within S1 sustain whole-brain resting-state rhythmicity

Which synapses inside S1 carry its contribution? We sorted the synapses with both endpoints in the same source cluster into four classes by polarity: excitatory-to-excitatory (E–E), excitatory-to-inhibitory (E–I), inhibitory-to-excitatory (I–E), and inhibitory-to-inhibitory (I–I) (Fig. 5a). By raw count, all four classes split across the clusters in nearly identical proportions (Fig. 5b), and S1’s internal composition matches that of S2 and S3 (Supplementary Fig. 11). Counting synapses does not distinguish S1. Weighting them by learned strength does: about half of the total E–I weight and half of the total I–E weight formed by the top-5% inhibitory neurons falls inside S1, against far less in S2 and S3 (Fig. 5c). S1 is where the strong heterotypic synapses of this essential inhibitory population sit, even though the underlying wiring composition is unremarkable.

**Figure 5.**
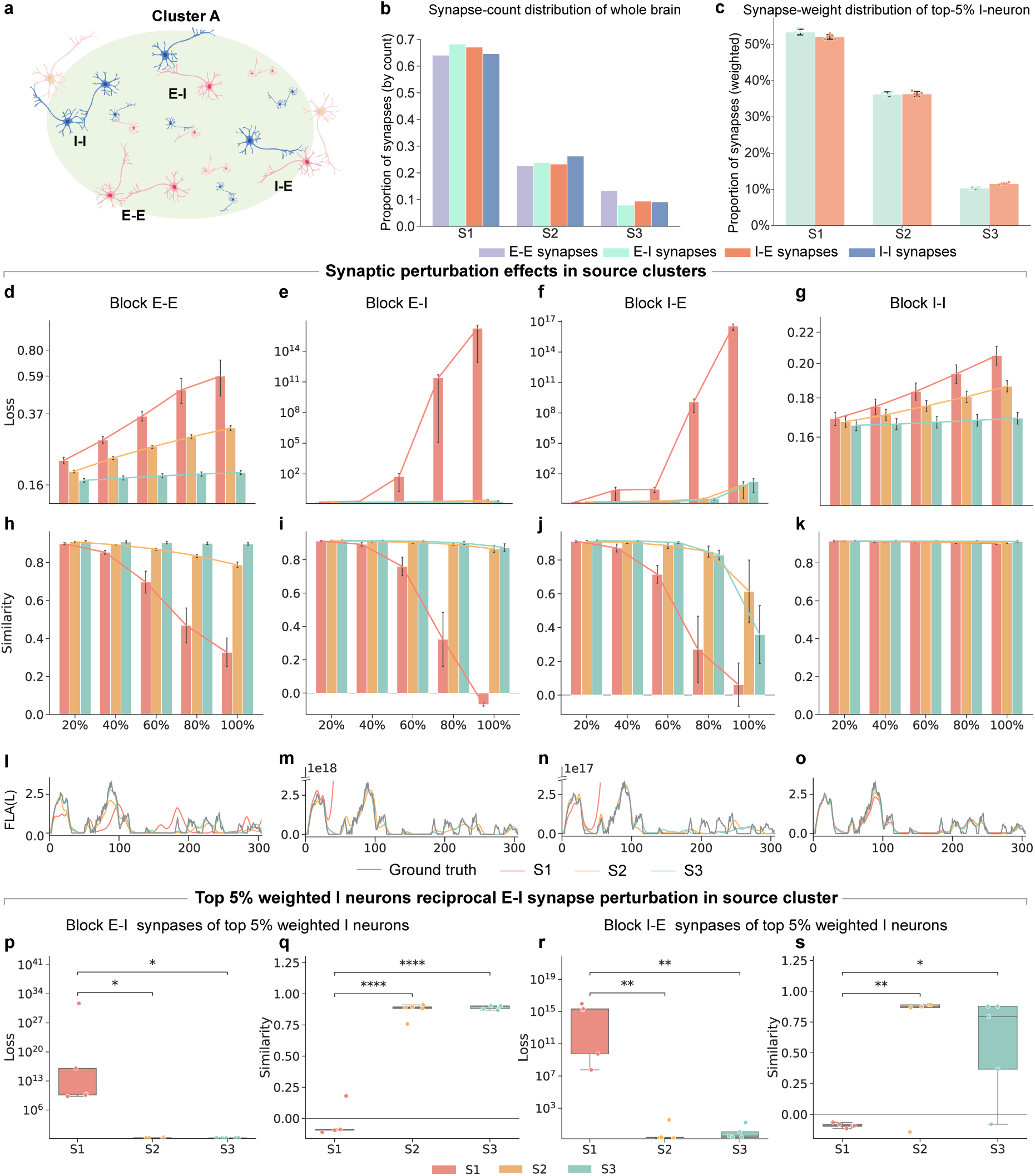
Reciprocal E–I pathways within source cluster 1 sustain whole-brain resting-state activity. **a**, Schematic of the four intra-cluster synaptic pathway types. Within each source cluster, connections between excitatory (E, blue) and inhibitory (I, red) neurons are classified into excitatory-to-excitatory (E–E), excitatory-to-inhibitory (E–I), inhibitory-to-excitatory (I–E), and inhibitory-to-inhibitory (I–I). **b,** Whole-brain cross-cluster distribution of intra-cluster synapses, by count, for each pathway type. Bars give the fraction of all whole-brain synapses of each type (E–E, E–I, I–E, I–I) that fall in S1, S2, or S3, the three source clusters defined in Fig. 3c; values sum to 1 across clusters within each type. All four pathway types show the same cross-cluster split, with ∼65% of synapses in S1, ∼22–26% in S2, and ∼8–13% in S3. **c,** Synaptic-weight–weighted cross-cluster distribution of the E–I (left) and I–E (right) synapses formed by the top-5% weighted inhibitory neurons identified in Fig. 4g. For each pathway type, the bar gives the fraction of total absolute synaptic weight Σ*_j→i_* |*w_ij_*| (rather than synapse count, as in **b**) that falls in S1, S2, or S3; values sum to 1 across clusters within each type. For both E–I and I–E, ∼52% of the total weight falls in S1, ∼36% in S2, and ∼10% in S3. Bars, mean ± std.; *n* = 5 trained models. **d–g,** Effect of progressive intra-cluster pathway blockade on test-set prediction loss. At each block ratio (20%–100%), synapses of the indicated pathway type within each source cluster were silenced and total loss was computed. **d,** E–E. **e,** E–I. **f,** I–E. **g,** I–I. Within S1, I–I blockade holds the loss within ∼0.16–0.22 across the full range. Logarithmic loss axes in **e** and **f**, where the simulated rates leave the bounded regime (Methods). Bars, mean ± std.; *n* = 5 independently trained models. **h–k,** Pearson correlation between simulated and empirical neuropil activity under the same progressive blockade as in **d–g**: **h,** E–E; **i,** E–I; **j,** I–E; **k,** I–I. Bars, mean ± std.; *n* = 5 trained models. **l–o,** Representative simulated activity traces of one example neuropil (FLA(L)) at 100% intra-cluster blockade of E–E (**l**), E–I (**m**), I–E (**n**), and I–I (**o**) pathways. Empirical ground truth in gray. **p–s,** Targeted blockade of the E–I incoming (**p**, loss; **q**, similarity) or I–E outgoing (**r**, loss; **s**, similarity) synapses of the top 5% weighted inhibitory neurons within each source cluster. Bars, mean ± std.; *n* = 5 trained models. Two-sided paired Student’s *t*-tests with Holm–Bonferroni correction: **p,** E–I loss: S1 vs S2, *p* = 1.02 × 10*^−^*^2^; S1 vs S3, *p* = 1.02 × 10*^−^*^2^. **q,** E–I similarity: S1 vs S2, *p* = 5.36 × 10*^−^*^5^; S1 vs S3, *p* = 4.63 × 10*^−^*^5^. **r,** I–E loss: S1 vs S2, *p* = 1.58 × 10*^−^*^3^; S1 vs S3, *p* = 1.07 × 10*^−^*^3^. **s,** I–E similarity: S1 vs S2, *p* = 9.12 × 10*^−^*^3^; S1 vs S3, *p* = 1.38 × 10*^−^*^2^. (The two **p** comparisons agree to three significant figures by coincidence; raw *p* = 1.02111 × 10*^−^*^2^ for S1 vs S2 and 1.01505 × 10*^−^*^2^ for S1 vs S3.) n.s. indicates *p* ≥ 0.05; exact corrected *p* values are reported above. Neuropil abbreviations as in Methods.

Blocking each class in turn, from 20% to 100% of its intra-cluster synapses, separates the four into two regimes. Intra-S1 E–E blockade raised loss from ∼0.16 to ∼0.85 while keeping rates bounded (Fig. 5d), and I–I blockade barely moved it (Fig. 5g). Correlation with the recordings stayed near baseline for both (Fig. 5h,k), and the example trace tracked the empirical envelope even at full blockade (Fig. 5l,o). E–I and I–E blockade behaved differently. Rates left the bounded regime (Fig. 5e,f), correlation fell to zero or below (Fig. 5i,j), and the trace departed from the recordings (Fig. 5m,n). The collapse built up gradually across the blockade range rather than at a single threshold, so the heterotypic loop carries no spare capacity. The same blockade inside S2 or S3 left both quantities at baseline in all four classes (Fig. 5d–k). E–I and I–E coupling matters only inside S1.

Can we localize this to the cells identified earlier? We silenced, cluster by cluster, the E–I inputs to or the I–E outputs from the top-5% weighted inhibitory neurons, leaving every other synapse in the brain intact (Methods). This touches the synapses of only ∼5% of inhibitory neurons, far fewer than the class-wide blockade. Inside S1 it still collapsed the model: loss rose by several orders of magnitude (Fig. 5p,r) and correlation fell to zero or below (Fig. 5q,s), with S1 differing from both S2 and S3 in every panel. Inside S2 or S3 the same intervention did nothing (Fig. 5p–s). These synapses are non-redundantly required, and the surrounding circuitry does not compensate for their loss.

### Per-cell-type silencing identifies a sparse, brain-spanning essential set that dissociates from cell-type enrichment

We next asked which cell types these neurons belong to. Mapping each FlyWire identifier onto the systematic adult-brain cell-type annotation^[7]^ lets us put two different questions to any population: which cell types are most abundant in it (Fig. 6a,c,e), and which, silenced one type at a time, collapse the model (Fig. 6b,d,f). We asked both of the shortened-timescale tail, of the top-5% weighted inhibitory neurons, and of the whole brain.

**Figure 6.**
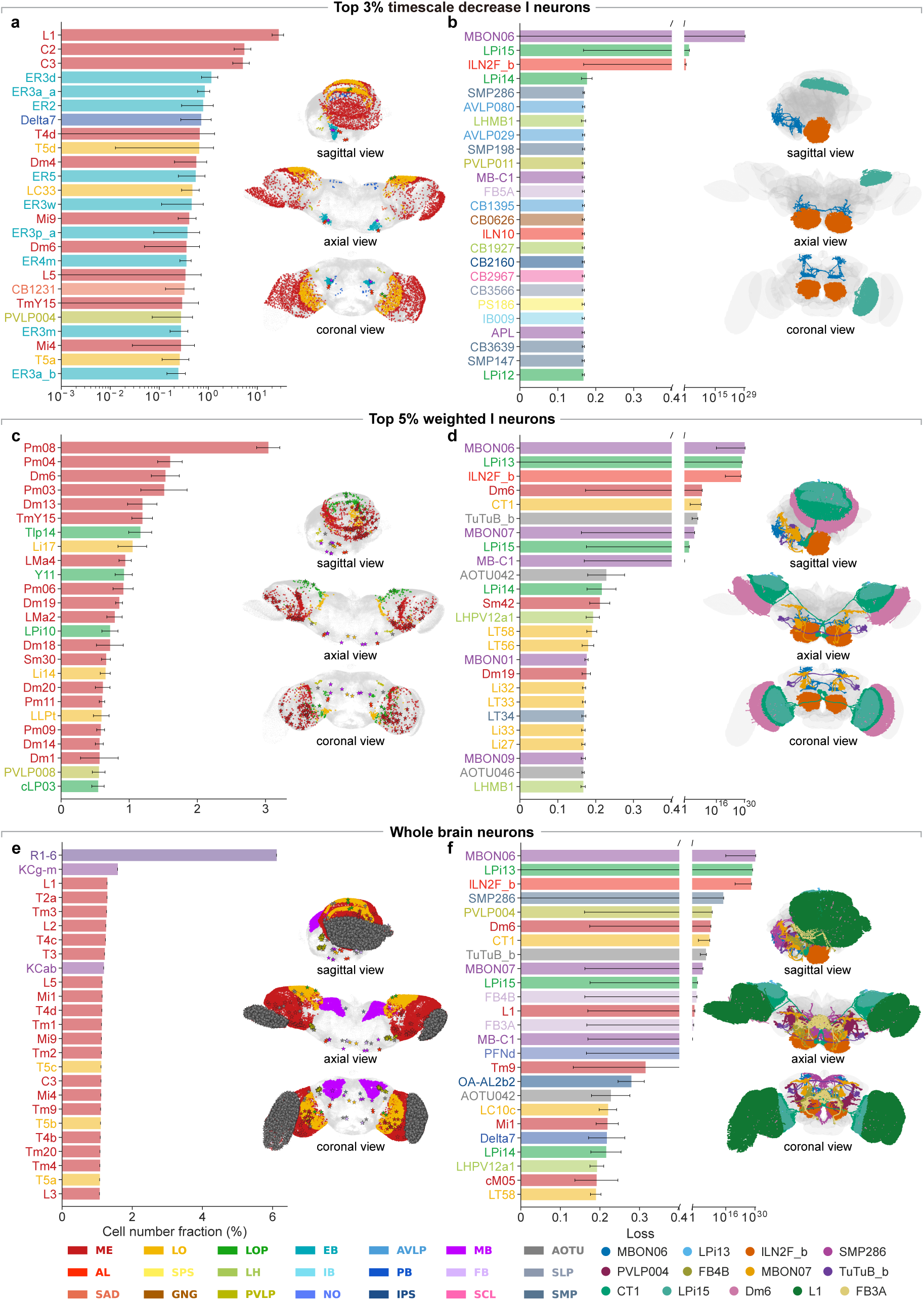
Cell-type enrichment and per-cell-type silencing dissociate, and identify a sparse brain-spanning essential set. **a**, Cell-type composition of the top 3% of inhibitory neurons with the most shortened intrinsic time constants (Δ*τ*-decrease tail of Fig. 4a,b), log scale. Each bar is the neuron count of that cell type within the top-3% population divided by the population’s total neuron count. Right, every cell type in the top-3% population mapped onto the standard *Drosophila* brain template (sagittal, axial, frontal views): each cell type is plotted as colored points at the soma positions of its member neurons, and pentagrams mark the cell types whose per-cell-type silencing loss in **b** exceeds 10. Cell-type labels are color-coded by primary neuropil of innervation (legend at bottom; neuropil abbreviations follow the Ito atlas^[15]^; cell-type names follow the FlyWire annotation^[7]^). Bars, mean across *n* = 5 trained models; individual model values as dots; whiskers, std. **b,** Per-cell-type silencing loss inside the top-3% population. For each cell type, all of its member neurons inside the population are set to silent and the model is re-run; the bar gives the resulting prediction loss against empirical activity. Top loss values are shown on a broken axis (inset to log scale). Right, full-arbor morphology of every cell type whose silencing loss in this panel exceeds 10, rendered on the standard brain template. Same color code and sampling as **a**. **c,d,** As in **a,b** but for the top 5% weighted inhibitory neurons (Fig. 4g–l); composition shown on linear scale. **e,f,** As in **a,b** but at the whole-brain level: **e** gives the brain-wide neuron-count fraction of each cell type (linear scale) together with the brain map of every cell type in the FlyWire annotation^[7]^, and **f** gives the per-cell-type silencing loss when every brain-wide member of the cell type is silenced, together with the morphology of those cell types whose loss exceeds 10. Only the 30–50 leading cell types per bar panel are shown; remaining types fell below the cutoff. Fractions in **a,c** are within-panel (each cell type’s value is its neuron count divided by the total neuron count of the selected population); fractions in **e** are brain-wide. Leading members in **a**: the lamina monopolar cells L1 (∼28% of the population) and L5, the centrifugal cells C2 and C3 (∼5% each), the motion detectors T4 and T5 with their medulla inputs Mi9 and Tm5c, and the ellipsoid-body ring neurons ER2, ER3a, ER3d, ER3m, ER4m and ER5 together with Delta7. Leading members in **c**: the Pm amacrines Pm03, Pm04, Pm06, Pm08, Pm09 and Pm11; the distal-medulla cells Dm6, Dm13, Dm14 and Dm18–Dm20; the lobula intrinsic neurons Li10–Li22; the lobula multicolumnar amacrines LMa2–LMa5; and the translobula-plate and lobula-plate intrinsic cells Tlp4, Tlp14 and LPi10. Neuron counts of the cell types whose silencing loss exceeds 10: L1, 1591; Dm6, 65; ILN2F_b, FB3A and MBON07, four each; MBON06, LPi13, LPi15, CT1 and TuTuB_b, two each^[7]^.

Training shortened intrinsic timescales in the fly’s experimentally fastest visual neurons (Fig. 6a). The lamina monopolar cell L1, the fastest documented relay in the fly visual system^[32, 33]^, made up ∼28% of this tail, roughly two-thirds of all L1 cells in the brain. Other early-visual types with brief integration windows followed: lamina cell L5, the centrifugal cells C2 and C3^[34, 35]^, and the motion detectors T4 and T5 with their medulla inputs Mi9 and Tm5c^[33, 36]^. A second, non-visual group comprised the ellipsoid-body ring neurons and Delta7, whose tens-of-milliseconds dynamics sustain the heading-direction ring attractor^[37, 38]^. The time constant was a free parameter, yet it converged onto the neurons with the shortest documented temporal kernels.

Silencing inside the same population returned a different set (Fig. 6b). Three cell types drove the loss to unbounded values: MBON06, a glutamatergic mushroom-body output neuron of the *β*1 compartment^[39]^; LPi15, a lobula-plate intrinsic neuron that gates direction selectivity^[40]^; and ILN2F_b, a GABAergic antennal-lobe interneuron^[41]^. All three are numerically rare, two to four cells each^[7]^. Abundance and indispensability come apart.

The top-5% weighted inhibitory hubs are mostly GABAergic local interneurons of the optic lobe (Fig. 6c). Pm amacrines predominate, mediating cross-columnar divisive normalization and contrast gain control^[42, 43]^; distal-medulla cells, lobula intrinsic neurons, lobula multicolumnar amacrines, and translobula-plate and lobula-plate intrinsic cells that supply opponent inhibition for direction selectivity make up the rest^[40]^. All five families show cell-type-specific patterned spontaneous activity from development into adulthood^[26, 28]^. Silencing inside the hub (Fig. 6d) again put MBON06 and ILN2F_b on top, followed by Dm6 (65 cells, also abundant in the hub^[34, 35]^) and a set of types with two to four cells each: CT1, the single giant amacrine spanning medulla and lobula and the optic lobe’s dominant non-Pm GABAergic source^[26, 42]^; TuTuB_b; MBON07; and LPi13 and LPi15. The hub is numerically dominated by optic-lobe interneurons, but the cell types that break it reach into the mushroom body, the central complex, and the columnar visual pathway.

Across the whole brain, raw counts are dominated by the chromatic photoreceptors R1–6, the *γ*-main Kenyon cells, the columnar projection neurons T2a, and the lamina monopolar cell L2 (Fig. 6e)^[7, 39]^. Silencing singled out none of these except L1 (Fig. 6f). The largest losses came instead from cell types spread across the optic lobe (CT1, Dm6, LPi13, LPi15, L1), the mushroom body (MBON06, MBON07), the central complex (FB4B, TuTuB_b, FB3A), the antennal lobe (ILN2F_b), and the higher-order protocerebrum (SMP286, PVLP004). All but Dm6 and L1 number four cells or fewer^[7]^. The same small set tops the ranking in all three populations, although the three were selected on different criteria. Silencing therefore selects on functional indispensability, not on population membership.

Where in the circuit does this set act? Blocking S1-routed synapses one cell type at a time returns MBON06 and Dm6 on the input side and LPi13, CT1, and LPi15 on the output side as the largest-loss perturbations (Supplementary Fig. 12a,b). The same block on S2-routed or S3-routed synapses produced no comparable loss, every cell type staying two orders of magnitude below the largest S1 effects (Supplementary Fig. 12c–f). The reciprocal E–I/I–E loop of Figs. 3 and 5 therefore has a cell-type signature, and it is confined to S1.

### A whole-brain cell-type atlas resolves the essential hubs into reciprocal excitatory–inhibitory partnerships

To see how these cell types are wired, we placed each essential inhibitory hub at the center of its cell-type-resolved ego-network on the brain template (Fig. 7). The hubs are spread across the brain: Dm6, CT1, LPi13, and LPi15 in the optic lobe, PVLP004 in the ventrolateral protocerebrum, MBON06 in the mushroom body, and FB4B and FB3A in the fan-shaped body. Most were flagged by both selection criteria, the shortened timescale and the high outgoing weight. Each sits at the center of a converging fan of mostly excitatory partner cell types, with its synapses routed through the source cluster of its host neuropil. The essential set is not a loose collection of cells but one wiring template repeated across the optic lobe, the mushroom body, and the central complex: an inhibitory hub locked to a small group of excitatory partners.

**Figure 7.**
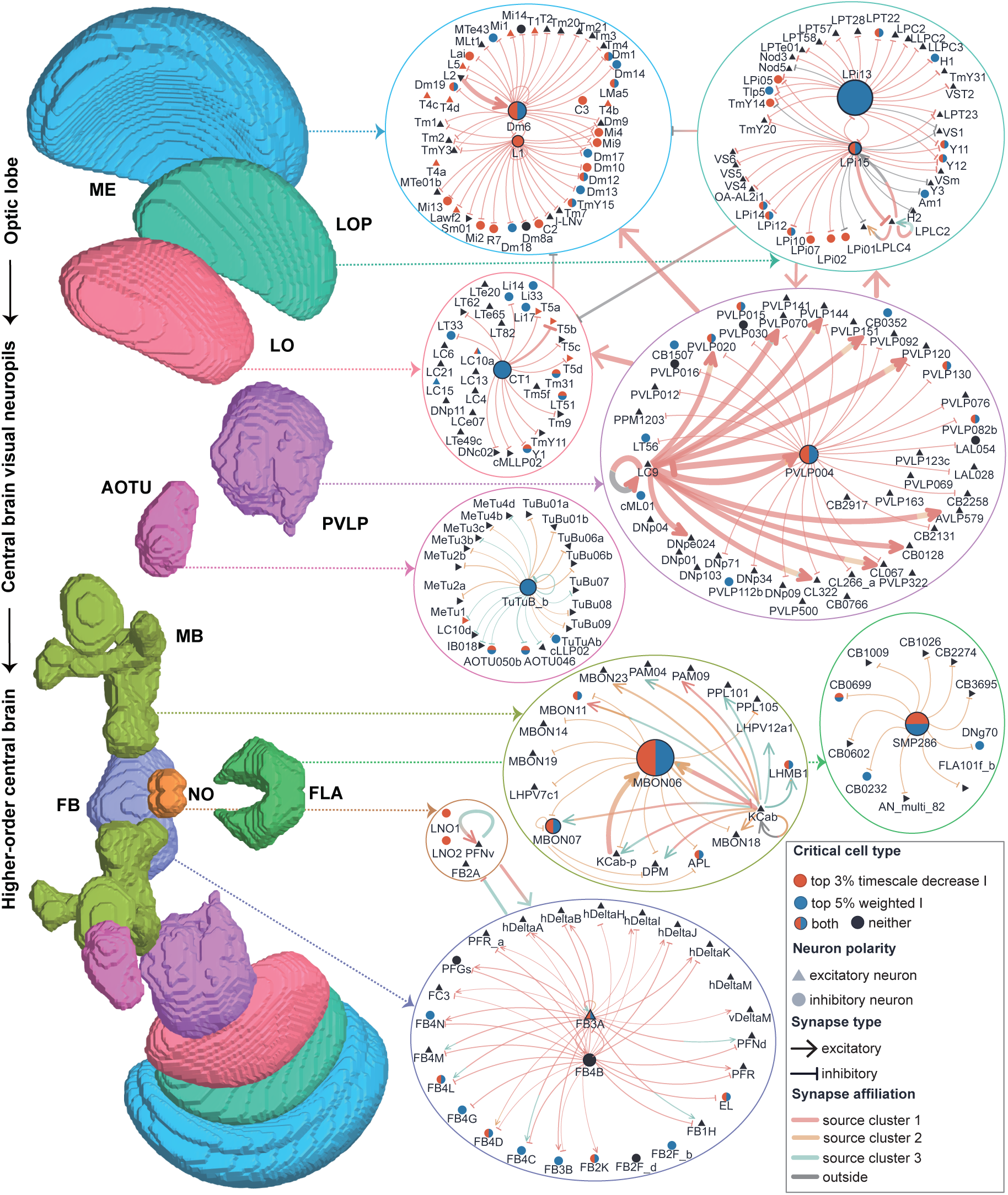
A whole-brain cell-type wiring atlas of the essential inhibitory hubs and their excitatory partners. Each essential inhibitory hub cell type returned by per-cell-type silencing (Fig. 6) is placed at the center of its cell-type-resolved ego-network, grouped by the neuropil that contains it and keyed to the standard *Drosophila* brain template (left; 3D renderings of ME, LO, LOP, AOTU, PVLP, MB, FB, NO, FLA). Nodes are cell types, and node size scales with the loss produced by silencing that cell type *in silico*, so the essential hub is the largest node in each network. Node shape gives neuron polarity (triangle, excitatory; circle, inhibitory). The fill color of each hub marks the criterion that selected it (red, top-3% shortened-timescale inhibitory population of Fig. 4a; blue, top-5% weighted inhibitory population of Fig. 4g; split red/blue, selected by both; black, neither). Edges are synapses, and edge width scales with the loss produced by blocking that connection *in silico*. The edge terminator gives synapse polarity (arrowhead, excitatory; bar, inhibitory). Edge color gives the source cluster through which the synapse is routed (blue, S1; orange, S2; green, S3; gray, outside; clusters as in Fig. 3c). The hubs span the optic lobe (Dm6 in ME, CT1 in LO, LPi13 and LPi15 in LOP), the optic glomeruli and ventrolateral protocerebrum (TuTuB_b in AOTU, PVLP004 in PVLP), the mushroom body (MBON06), the central complex (FB4B and FB3A in FB, with a PFNv–FB2A– PFR_a chain in NO), and a flange network centered on SMP286. In most networks the central hub is an inhibitory cell flagged by the timescale or weight criterion of Fig. 4, and its converging partners are predominantly excitatory. The synapses route through the source cluster of the host neuropil: the S1-resident hubs (Dm6, CT1, LPi13/LPi15, PVLP004, MBON06, FB4B/FB3A) are dominated by S1 (blue) synapses, the AOTU and flange hubs (TuTuB_b, SMP286) by S2 (orange), and the noduli chain by S3 (green). The reciprocal excitatory–inhibitory motif of Figs. 3–5 therefore recurs at single-cell-type resolution and across the brain. Cell-type names follow the FlyWire annotation^[7]^; neuropil abbreviations follow the Ito atlas^[15]^.

If that is the architecture, a hub and its partners should stand or fall together. They do (Supplementary Fig. 13). Silencing an inhibitory hub alone drove the loss to unbounded values. Silencing its principal excitatory partner alone left the model at baseline. Silencing both together returned it to baseline. This held for five pairs: PVLP004 with LC9, FB4B with PFNv, MBON06 with the Kenyon cells KCab and KCab-p, LPi15 with LPLC2, and Dm6 with L2 (Supplementary Fig. 13d–h). CT1 moved the same way, the combination lowering the loss by several orders of magnitude though not fully to baseline (Supplementary Fig. 13i). The co-dependence also held at the synapse level: blocking the hub’s inhibitory projection onto its partner collapsed the model, and additionally blocking that partner’s downstream excitatory projections restored baseline activity (Supplementary Fig. 13j–m). The instability that follows silencing an essential inhibitory cell is therefore disinhibition. Removing the inhibition releases one specific excitatory partner into runaway activity; removing that partner, as a cell or as a projection, absorbs it again. Ranking synapses by the loss their removal causes names the same partnerships: excitatory inputs onto a hub (KCab and KCab-p onto MBON06, L2 onto Dm6) and inhibitory outputs from a hub onto its excitatory targets (PVLP004 onto LC9, FB4B onto PFNv, LPi15 onto LPLC2, CT1 onto T5b), and the same partnerships top the neuropil-pair ranking among optic-lobe and central-complex regions (Supplementary Fig. 13a–c). The unit of indispensability is the reciprocal partnership, not the inhibitory cell alone.

## Discussion

Connectome-constrained whole-brain modeling can do more than reproduce neural activity: it can name the cells and synapses that generate it. Fitting a cell-resolution network constrained by the adult *Drosophila* connectome to whole-brain calcium recordings produced a model that reproduced signatures it was never trained on, lognormal synaptic weights and scale-free neuronal avalanches. Its functional connectivity generalized to held-out data only when the anatomical wiring, the readout geometry, and the recurrent topology were kept; expressive capacity alone did not substitute.

Perturbing this model localized the resting-state scaffold to a compact core of 14 source neuropils, an axis running from fast sensory relays in the optic lobe to the mushroom body and the central complex. Inside that core the causal weight rests on a sparse population of high-weight inhibitory hubs and their reciprocal E–I loops, which carry no redundancy. At cell-type resolution the same wiring blueprint recurs across the sensory-to-central hierarchy: removing an inhibitory cell destabilizes brain-wide activity by releasing one specific excitatory partner into runaway disinhibition. Resting-state activity in this model is therefore organized by a sparse, connectome-defined circuit rather than by uniformly distributed interactions, and the slow, coherent, near-critical dynamics reported across species acquire a candidate cellular and synaptic substrate.

Connectomes and brain-wide recordings have lived in separate workflows because nothing has forced them to share parameters. Pure simulation^[44]^ propagates activity through a fixed connectome but cannot tune effective weights against recordings. Task-driven models^[45]^ recover weights sufficient for a behavioral objective, not weights consistent with the activity neurons actually exhibit. Mean-field approaches such as The Virtual Brain^[46]^ match higher-order statistics of population activity but, by construction, cannot name a neuron or a synapse. Fitting the connectome to recordings inverts the arrangement: the recordings dictate *what* the model reproduces, the connectome dictates *how*. Activity is the loss function, the connectome the substrate that realizes it, and mechanistic claims at single-cell and single-synapse resolution follow from two datasets that alone cannot deliver them. The fly is one worked example of a recipe. Any pairing of a synapse-resolution connectome with concurrent or matched whole-brain recordings^[3, 47]^ can be fit the same way, yielding an *in silico* preparation in which the effect of any cell or synapse on global dynamics can be measured directly.

Several limitations bound this interpretation. The neuropil clusters are bilaterally asymmetric. Asymmetries in LOP, AVLP and PVLP partly reflect how the recordings were acquired^[9]^, whereas those in MB-ML and PLP may be optimization artifacts and need bilateral recordings to resolve. FlyWire also omits gap junctions^[48]^, graded transmission^[49]^, and neuromodulation^[50, 51]^, all of which shape whole-brain computation. The cellular and synaptic mechanisms identified here therefore remain model-derived hypotheses that require *in vivo* validation.

## Methods

### Experimental whole-brain calcium imaging data

Whole-brain resting-state calcium imaging of head-fixed adult *Drosophila melanogaster* expressing the pan-neuronal calcium indicator GCaMP6s was taken from Turner et al.^[9]^. Each volume was acquired at 1.2 Hz, motion-corrected, and registered to the standard fly brain template. Activity was averaged within each of the 73 anatomically defined neuropils of the Ito reference atlas^[15]^ to produce neuropil-level Δ*F/F* traces. The resting state was defined as the absence of locomotion and prominent grooming throughout the recording, following the criteria of the original study. Each fly in the source dataset contributed at least 14 minutes of imaging. All analyses are performed on an example fly, with at least five independently trained model instances per condition (*n* = 5 trained models throughout); the first half of that recording served as the training set, and the second half was held out for all generalization, effectome, and ablation analyses reported in the main text.

### Calcium-to-firing-rate deconvolution

The model generates per-neuron firing rates, while the measurements are slow, neuropil-averaged Δ*F/F* traces. To train against a signal the model can reproduce, we inverted the calcium forward model and recovered, for each neuropil, an instantaneous estimate of the local population firing rate, not individual spike trains. A neuropil Δ*F/F* trace averages thousands of single-neuron signals, which linearises the relationship between fluorescence and underlying activity; the natural latent variable is therefore the continuous non-negative *mean firing rate* of the neuropil, not a binary spike sequence^[52, 53]^.

### Forward model

For each neuropil *k*, the measured calcium trace *c_k_*(*t*) was modeled as a noisy linear convolution of the underlying population firing rate *F_k_*(*t*) ≥ 0 with the indicator impulse response *h*(*t*),

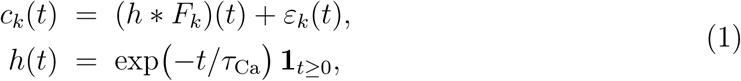

where *τ*_Ca_ = 1.5 s is the GCaMP6s decay time constant^[54]^; the rise time falls well below the volumetric sampling interval Δ*t* = 1*/*1.2 s and is therefore unresolved by the imaging, so *h* is written as a pure decay. *ε_k_*(*t*) is i.i.d. Gaussian measurement noise. Equation (1) is exact in the limit of (i) linearity of Δ*F/F* in instantaneous rate at the single-neuron level and (ii) spatial averaging over many neurons within a neuropil. Both conditions hold for the 73 anatomically large neuropils analyzed here.

### Recovery as a non-negative regularized inverse

Inverting Eq. (1) is ill posed: *h* acts as a low-pass filter, so high-frequency content of *F_k_* is unrecoverable from *c_k_* alone. We resolved this by imposing two priors that follow from the physics of the signal. *F_k_* is non-negative. *F_k_* varies smoothly on the imaging timescale because it averages over many independent neurons. We then estimated *F*^^^*_k_* as the constrained Tikhonov-regularized least-squares solution^[55]^

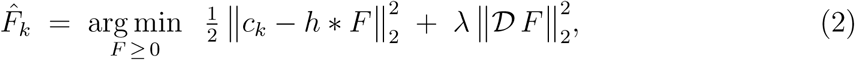

with D the first-order temporal-difference operator and *λ >* 0 a smoothness weight. The objective is convex; we minimized it with FISTA^[56]^ composed with projection onto the non-negative orthant. The regularization strength *λ* was selected per neuropil by the L-curve criterion^[55]^; the selected values clustered around *λ* ≈ 10*^−^*^1^. After convergence, the deconvolved trace *F*^^^*_k_* (*t*) was rescaled by a single global multiplicative factor so that its mean across all neuropils and all time bins equalled 1 Hz, giving units interpretable as a population firing rate. *F*^^^*_k_* (*t*) is used throughout as the empirical training and testing target, and is directly comparable to the simulated neuropil rate *R_k_*(*t*) of Eq. (6).

### Connectome data and neuron–synapse assignment

The structural scaffold came from FlyWire v783, a synaptic-resolution connectome of the adult *Drosophila* brain^[1, 7]^. After standard quality filtering, the model included 138,639 neurons and 15,091,983 chemical synapses. Each model neuron was mapped one-to-one onto a FlyWire neuron and inherited its presynaptic partners, postsynaptic targets, synapse counts, and excitatory/inhibitory polarity prediction. Polarity labels were taken from FlyWire neurotransmitter predictions^[7]^ and were frozen during training; the whole-brain excitatory fraction was approximately 0.6. The number of synapses each neuron *i* formed within neuropil *k, n^syn^_i,k_*, was extracted from the FlyWire synapse table and used for both the neuropil-level readout (below) and the cluster-membership assignments. The connectome scaffold and Eq. (3) jointly omit several biological elements: gap-junctional and graded transmission, which are not reconstructed in FlyWire v783; neuropeptidergic signaling; and neuromodulatory state, which is folded into the fixed effective weights {|*w_ij_*|} rather than modeled as a separate dynamical variable. These omissions are addressed in the Discussion.

### Neuropil nomenclature and abbreviations

Neuropils are referred to throughout the manuscript, figures, and Supplementary Table 1 by the standard three- to four-letter abbreviations of the Ito et al. reference atlas^[15]^, with suffixes (L), (R), and (L/R) denoting the left hemisphere, the right hemisphere, and bilateral pairs, respectively; cell types follow the FlyWire adult-brain annotation^[7]^. Full abbreviation-to-name mappings are given in Supplementary Table 2 (neuropils, grouped by anatomical superregion) and Supplementary Table 3 (cell types).

### Single-neuron dynamics and whole-brain network construction

Each of the 138,639 neurons followed a first-order firing-rate equation,

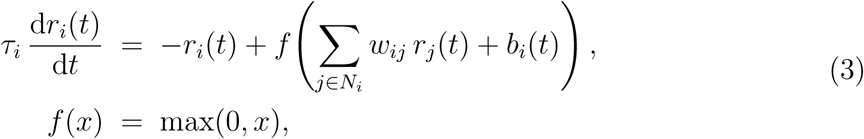

where *r_i_*(*t*) is the instantaneous firing rate of neuron *i*, *τ_i_* its intrinsic time constant, *N_i_* the set of presynaptic partners specified by the FlyWire connectome, and *f* a rectified-linear activation that constrains rates to be non-negative. The effective synaptic weight from presynaptic neuron *j* to postsynaptic neuron *i* factorises as *w_ij_* = sgn(*j*) |*w_ij_*|, with the connectome-derived polarity sgn(*j*) ∈ {+1, −1} frozen during training and only the magnitude |*w_ij_*| learned. The stochastic drive

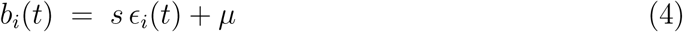

combines a scalar global noise scale *s* and offset *µ*, shared across the population, with independent standard Gaussian noise *ɛ_i_*(*t*) ∼ N(0, 1) resampled separately for each neuron at each integration step. Equation (3) was integrated with the exponential Euler scheme at Δ*t* = 1*/*1.2 s, matching the volumetric imaging rate, yielding the discrete-time update used during simulation and training,

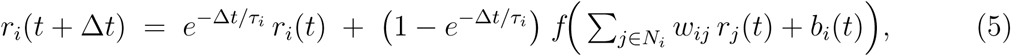

with *b_i_*(*t*) as in Eq. (4). Equation (5) is the form through which gradients are propagated (see below); the exponential Euler scheme integrates the linear −*r_i_/τ_i_* term exactly, so the discrete update is stable even when *τ_i_* approaches Δ*t*. The trainable parameter set was the synaptic-weight magnitudes {|*w_ij_*|} (≈ 1.5 × 10^7^), the per-neuron time constants {*τ_i_*}, and the two global drive parameters (*s, µ*).

### Connectome-derived neuropil readout

To compare simulated single-neuron activity with neuropil-resolved empirical recordings, single-neuron rates were projected onto a 73-dimensional neuropil space through a synapse-density-weighted readout. The simulated activity of neuropil *k* at time *t* was

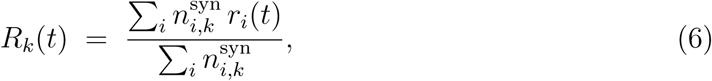

i.e., a normalized average of single-neuron rates weighted by the number of synapses each neuron forms within neuropil *k*. The readout has no learnable parameters of its own; it is fully specified by the FlyWire synapse counts and imposes a strict anatomical bias on the model. *R_k_*(*t*) served as the model-side signal in the training loss and as the basis for all downstream neuropil-level analyses. As a control (Fig. 2i,j), the readout was replaced by a parameter-matched unconstrained linear projection with freely learnable weights initialized from a small-variance Gaussian.

### Training data, loss function, and optimization

Free parameters were estimated by gradient descent to minimize the discrepancy between simulated and empirical neuropil activity. The loss was the time-averaged mean squared error across all 73 neuropils and all training or held-out timepoints,

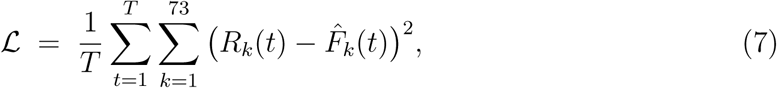

with *R_k_*(*t*) the simulated neuropil rate (Eq. (6)) and *F*^^^*_k_* (*t*) the deconvolved empirical firing-rate proxy (Eq. (2)). No clipping or saturation was applied at any stage; the loss values exceeding 10^16^ reported in Fig. 3e,f, Fig. 4h,j,l and Fig. 5e,f,p,r reflect numerical divergence of the rate dynamics under perturbations that break excitation–inhibition balance, and are plotted on a logarithmic axis. Their magnitude reflects how far the integrator advanced before overflow, not the size of an effect, so differences among these divergent values are not meaningful. Gradients through the recurrent dynamics of the 138,639-neuron, 15,091,983-synapse network were computed by backpropagation through time^[57]^, with gradient checkpointing bounding the peak memory of long rollouts. As an alternative training mode, the same models were trained with BrainTrace^[58]^, an online gradient propagator that pushes gradients along the simulated trajectory without materializing the full backward graph. The online scheme gave fits of comparable quality to BPTT and confirmed that the results do not depend on the gradient estimator. Parameters were updated with Adam^[59]^ (learning rate 10*^−^*^3^; default momentum parameters *β*_1_ = 0.9 and *β*_2_ = 0.999; batch size 64). Gradient optimization was applied over the full training segment without truncation.

The per-neuron initial state *r_i_*(0) was a free parameter learned jointly with the synaptic weights, time constants, and global drive parameters. At test time, the model was first rolled forward from this learned initial state over the training segment, and then continued autoregressively over the held-out test segment under the same stochastic-noise regime as training; all reported predictive metrics (loss, Pearson correlation, functional connectivity, avalanche statistics) were computed on the test-segment trajectory, averaged across *n* = 5 independent noise seeds per condition.

For each fly we trained at least five independent model instances from different random seeds. Two priors over the initial weight magnitudes were tested: a uniform interval and a truncated-normal prior centered at zero. Both converged to statistically indistinguishable weight distributions (Fig. 2o,p). Time constants *τ_i_* were initialized by independent draws from a uniform distribution and refined during training. No separate validation segment was held out; training terminated when the training loss plateaued (Fig. 1f).

### Baseline and control architectures

To isolate the contribution of the connectome scaffold, we compared the connectome-constrained model against two families of parameter-matched generic baselines that carried no connectome structure: a gated recurrent unit (GRU) as a recurrent generative model, and multilayer perceptrons (MLPs) trained under the next-step prediction paradigm. Both shared the observables (the 73-dimensional neuropil rates), the train/test split, the MSE loss (Eq. (7)), and the Adam^[59]^ schedule of the connectome-constrained model, and each was matched to it in total parameter count to within ±5% (≈ 1.5 × 10^7^). The two MLP variants of Fig. 2k,l differ only in the readout: a freely learned linear projection versus the fixed connectome-derived readout of Eq. (6). Full architecture, training, and evaluation details for both baselines are given in Supplementary Note 1.

### Software, hardware, and computational resources

All whole-brain models were built, simulated, trained, and analyzed within the BrainX ecosystem (version 2026.3.12)^[58, 60–62]^, a JAX-compatible runtime for large-scale neural-network simulation with physical-units enforcement and the online optimization system used here. Training and *in silico* perturbation experiments ran on a single NVIDIA RTX 6000 Pro GPU; per-experiment wall-clock times are documented in the released training scripts (see Code availability). Random seeds for parameter initialization, noise injection, and resampling controls were fixed and logged for every experiment to support reproduction.

### Avalanche detection and power-law fitting

Neuronal avalanches were detected on a common binary event raster obtained by thresholding each continuous activity trace at three times its baseline standard deviation^[14, 63]^, applied identically to the deconvolved empirical neuropil rate *F*^^^*_k_* (*t*) (Eq. (2); Fig. 1i), the simulated neuropil rate *R_k_*(*t*) (Eq. (6); Fig. 1j), and the simulated single-neuron rate *r_i_*(*t*) (Eq. (3); Fig. 1k), with bin width Δ*t*_aval_ = 1*/*1.2 s on the held-out test segment. Avalanche size and duration distributions were fit with truncated power laws by maximum likelihood^[64]^, the same framework applied to the trained synaptic-weight magnitudes (Fig. 1h), and the truncated power law was compared against lognormal and exponential alternatives by Vuong-corrected likelihood-ratio tests with the powerlaw package^[65]^. The operational definition of criticality, the event-detection and avalanche definitions, and the full fitting and model-comparison procedure are given in Supplementary Note 2; the interpretation of the likelihood-ratio *p* values at the large sample sizes used here is discussed in Supplementary Note 3.

### Neuropil effectome construction and clustering

To map directed causal interactions between neuropils, for each source neuropil *s* we ablated every chemical synapse whose FlyWire-annotated terminal lies in *s* and measured the resulting change in prediction loss at every target neuropil. For a source-neuropil set *S* ⊆ {1*, …,* 73}, the perturbed weight matrix was

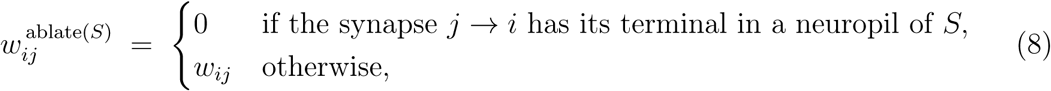

applied only to the recurrent term of Eq. (3); time constants *τ_i_*, polarities sgn(*j*), and the intrinsic drive *b_i_*(*t*) were left untouched. The synapse-level neuropil label was taken directly from the FlyWire v783 synapse table^[1, 7]^, so no neuron-to-neuropil assignment was required; a neuron whose synaptic terminals span multiple neuropils contributes to the ablation in each of those neuropils independently. For each ablation we simulated the perturbed model on the held-out test segment with five independent noise seeds and computed the per-target-neuropil loss change

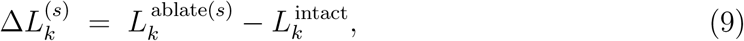

populating a 73 × 73 directed loss-change matrix indexed by source *s* (row) and target *k* (column). The effectome matrix shown in Fig. 3b is the mean of these per-model matrices across the *n* = 5 independently trained model instances; clustering was performed on this averaged matrix, and the resulting source-cluster (S1, S2, S3) and target-cluster (T1, T2, T3) assignments were then applied to each individual trained model in all subsequent cluster-based analyses.

Source and target neuropils were clustered independently by hierarchical agglomerative clustering applied to the row vectors (for sources) and column vectors (for targets) of the effectome matrix, with Euclidean distance and average linkage. The number of clusters was selected by maximizing the silhouette coefficient over *k* ∈ {2*, …,* 12}; this returned *k* = 3 source clusters (S1, S2, S3) and *k* = 3 target clusters. The silhouette curves for the source and target clusterings are shown in Supplementary Fig. 5; per-model reproducibility of the *k* = 3 source and target partitions, quantified by silhouette coefficient and Adjusted Rand Index against the averaged clustering, is reported in Supplementary Fig. 6. The dendrograms in the margins of Fig. 3b correspond to this clustering. Cluster assignments were projected onto the standard *Drosophila* brain template using the centroid coordinates of each neuropil.

For the selective-preservation experiments of Fig. 3d–g, cluster preservation meant ablating, through Eq. (8), every chemical synapse whose terminal lies outside the union of neuropils of the preserved cluster set, with *S* set to the complement of the preserved neuropils. Prediction loss and Pearson correlation between simulated and empirical neuropil activity were computed on the held-out test segment with five independent noise seeds per condition.

### Subpopulation identification and *in silico* silencing

For each neuron we computed the training-induced change in intrinsic time constant, Δ_τi_ = τ^post^_i_ − τ^init^_i_, averaged across independently trained model instances. Two tail subpopulations were taken from the extremes of this distribution: the 3% with the largest negative Δ*τ* (“decrease tail”) and the 3% with the largest positive Δ*τ* (“increase tail”) (Fig. 4a). To map their anatomical layout (Supplementary Fig. 10), the soma coordinates of each FlyWire neuron were registered to the standard brain template and rendered hemisphere by hemisphere.

Neuron silencing was implemented by clamping *r_i_*(*t*) ≡ 0 for every neuron *i* in the silenced set throughout the test simulation; this is distinct from the outgoing-weight ablation operator used for the effectome. Block-ratio silencing (Fig. 4b,e,f) cumulatively silenced the top *x*% of neurons ranked by Δ*τ* within each tail, sweeping *x* ∈ {0.5, 1, 2, 3, 4, 5}%. Random-control sets were drawn by sampling the same number of neurons uniformly without replacement; ten random draws were generated per condition. Cell-type-restricted silencing (Fig. 4e,f) used the same percentile ranking and block-ratio sweep but drew only from the excitatory or only from the inhibitory subpopulation.

A complementary essential-neuron set was defined directly from the learned synaptic weights. For each inhibitory neuron *i*, two scalar summaries were computed. The first, *W^in^_i_*, sums the absolute weights of synapses incoming to *i* (where *i* is on the postsynaptic side):

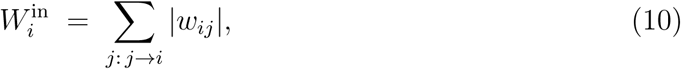

used in Fig. 4i,j. The second, *W^out^_i_*, sums the signed weights of synapses outgoing from *i* (where *i* is on the presynaptic side), so that for an inhibitory neuron *i* all terms share the negative polarity and *W^out^_i_* < 0:

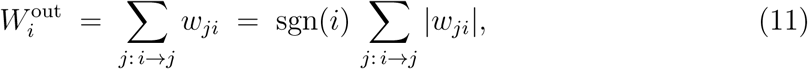

used in Fig. 4k,l. The essential inhibitory population is the top-5% of inhibitory neurons ranked by |*W^out^_i_*|: Fig. 4g plots the distribution of *W^out^_i_* across the inhibitory population (signed, negative for inhibitory neurons), and Fig. 4h shows the prediction loss when neurons in each percentile bin of |*W^out^_i_*| are silenced *in silico*. This single top-5% subset is used throughout Fig. 4i–l, Fig. 6c,d, and Supplementary Fig. 12; panels i, j of Fig. 4 analyze the incoming synapses of the subset (per-synapse aggregates of *W^in^_i_* across clusters), and panels k, l its outgoing synapses (per-synapse aggregates of *W^out^_i_* across clusters).

All cluster attribution and cluster-restricted blockade analyses on the essential inhibitory population (the top-5% subset defined above by |*W^out^_i_*|; Fig. 4g) used a per-synapse rule consistent with the effectome ablation of Eq. (8): a synapse was attributed to cluster *C* when its FlyWire-annotated terminal lay in a neuropil of *C*. With this convention, the source-cluster composition of Fig. 4i tallied every incoming (pre-synaptic) synapse of a top-5% neuron by the cluster containing its presynaptic terminal; the composition of Fig. 4k tallied every outgoing (post-synaptic) synapse of a top-5% neuron by the cluster containing its postsynaptic terminal; the fractions reported are sums of |*w_ij_*| over qualifying synapses divided by the across-cluster total. For the cluster-restricted blockade of Fig. 4j, only those incoming synapses to top-5% neurons whose presynaptic terminal lay in the targeted cluster were zeroed through Eq. (8); for Fig. 4l, only the corresponding outgoing synapses (postsynaptic terminal in the targeted cluster) were zeroed. For the cell-type tallies of Supplementary Fig. 12, an incoming synapse to a top-5% neuron was counted as routed through cluster *C* ∈ {S1, S2, S3} if its presynaptic terminal lay in a neuropil of *C*, and an outgoing synapse was counted as routed through *C* if its postsynaptic terminal lay in a neuropil of *C*; each qualifying synapse contributed its absolute weight |*w_ij_*| to the cell type of the corresponding partner (panels a, c for S1; e, g for S2; i, k for S3). The cluster-restricted per-cell-type block (Supplementary Fig. 12b, d, f, h, j, l) zeroed exactly the qualifying synapses of one cell-type/cluster/direction combination at a time and re-ran the model under the protocol of Fig. 5p,r.

### Intra-cluster pathway dissection

Intra-cluster synapses were classified into four pathway types by the polarities of their endpoints: E–E, E–I, I–E, and I–I (Fig. 5a). A synapse *j* → *i* was treated as intra-cluster for source cluster *C* when its FlyWire-annotated terminal lay in a neuropil of *C*, consistent with the per-synapse rule of Eq. (8). Polarity labels followed the connectome-derived signs used during training.

For Fig. 5b, the cross-cluster distribution of each pathway type was computed by counting intra-cluster synapses of that type in each source cluster and normalizing so that the three cluster values summed to one. For Fig. 5c, the same cross-cluster split was computed for the E–I and I–E synapses of the top-5% weighted inhibitory neurons of Fig. 4g, but with each synapse weighted by the absolute value of its learned weight |*w_ij_*| rather than counted once, so that the reported fractions are sums of |*w_ij_*| per cluster divided by the across-cluster total. Panel b therefore reflects synapse counts; panel c reflects synaptic weight. For Supplementary Fig. 11, the same intra-cluster synapse counts were instead normalized *within* each source cluster, so that the four pathway-class values summed to one per cluster, giving the within-cluster pathway-class composition of S1, S2, and S3.

For the progressive pathway-blockade experiments (Fig. 5d–o), at each block ratio *b* ∈ {20, 40, 60, 80, 100}% an independent uniformly random subset comprising *b*% of the intra-cluster synapses of the targeted pathway type was drawn afresh per repeat (no nesting across ratios) and its weights *w_ij_* set to zero (Eq. (8) restricted to the chosen synapse set). All other synapses were left intact. Prediction loss and Pearson correlation to empirical activity were computed on the held-out test segment with five independent simulation repeats per condition.

For the targeted-blockade experiments (Fig. 5p–s), the same procedure was applied to a fixed synapse set rather than a random subset: the E–I synapses incoming to, or the I–E synapses outgoing from, the top-5% weighted inhibitory neurons of Fig. 4g, restricted to each source cluster *C* in turn (i.e., to those synapses whose terminal lay in a neuropil of *C*).

### Cell-type assignment

Cell-type identities for the three subpopulations of Fig. 6a, c, e and for the partner cell types tallied in Supplementary Fig. 12 were obtained by joining each FlyWire neuron identifier against the systematic adult-brain cell-type annotation of Schlegel et al.^[7]^. Neurons with no annotated cell type were retained in the denominator but excluded from the per-type tallies. For Fig. 6a, c the denominator is the size of the subpopulation (top 3% by Δ*τ* decrease, and top 5% by outgoing weight magnitude |*W^out^_i_*| of Eq. (11), respectively); for Fig. 6e the denominator is the whole-brain neuron count. For Supplementary Fig. 12, the denominator of each weight-fraction panel is the total absolute synaptic weight |*w_ij_*| of cluster-routed synapses in the corresponding direction (incoming or outgoing) within the targeted cluster (*C* ∈ {S1, S2, S3}). Reported fractions are the mean across the *n* = 5 independently trained model instances, with individual model values shown as dots; only the 30–50 most frequent or most weight-contributing cell types per panel are displayed.

### Cell-type-pair silencing and the whole-brain cell-type atlas

The essential cell types analyzed in Fig. 6f and the cell-type atlas of Fig. 7 are the cell types whose whole-brain per-cell-type silencing (Fig. 6f) produced the largest increase in test-set prediction loss. The silencing operators below act on the recurrent term of Eq. (3) and leave the time constants *τ_i_*, polarities, and intrinsic drive unchanged; each readout is the test-set loss of Eq. (7), averaged over the *n* = 5 trained models with five noise seeds each.

For the per-pair projection silencing of Supplementary Fig. 13a,b, every chemical synapse of an ordered cell-type pair *A* → *B* (presynaptic neuron of type *A*, postsynaptic neuron of type *B*) was set to zero through Eq. (8), the model re-run, and the loss recorded. On the input side (Supplementary Fig. 13a) *B* was a essential cell type and *A* ranged over its presynaptic partner types; on the output side (Supplementary Fig. 13b) *A* was a essential cell type and *B* ranged over its postsynaptic partner types. For the neuropil-pair silencing of Supplementary Fig. 13c, synapses were grouped by the directed pair of neuropils containing their presynaptic and postsynaptic terminals and by polarity, each group was zeroed together, and the summed loss recorded and colored by the group’s polarity.

For the single-versus combined-cell-type silencing of Supplementary Fig. 13d–i, one or more whole cell types were silenced by clamping *r_i_*(*t*) ≡ 0 for all of their member neurons, as in the subpopulation silencing above, and three conditions were compared: the inhibitory hub alone, its excitatory partner cell type(s) alone, and the hub together with its partner(s). For the single-versus combined-projection silencing of Supplementary Fig. 13j–m, the inhibitory projection from a hub onto its excitatory partner was zeroed through Eq. (8), either alone or together with the excitatory projections from that partner onto its own downstream targets.

In the cell-type atlas of Fig. 7, each essential inhibitory hub is drawn at the center of an ego-network of its directly connected partner cell types, positioned by the neuropil that contains the hub and rendered on the standard brain template^[15]^. Node size encodes the per-cell-type silencing loss of that cell type and edge width the per-projection silencing loss of that connection, both as defined above; node shape encodes neuron polarity, hub fill the selection criterion of Fig. 4a,g, and edge color the source cluster (Fig. 3c) containing the synapse’s terminal.

### Statistical analysis

Sample sizes and the corresponding replication units are reported in each figure caption. Two replication units appear throughout: *n*_models_ ≥ 5 for independently trained model instances on the single example fly, and *n*_repeats_ = 5 for noise-seed repeats under a fixed model and perturbation condition. Error bars and shaded bands denote mean ± s.d. across the relevant replication unit unless stated otherwise.

Power-law and lognormal fits used the maximum-likelihood framework of Clauset et al.^[64]^; goodness of fit was reported as the Kolmogorov–Smirnov statistic, and competing distributions were compared by likelihood-ratio tests with the Vuong correction. Likelihood-ratio test *p* values are reported numerically and should be interpreted in the context of the large sample sizes involved (Supplementary Note 3). Cross-cluster comparisons of source-cluster composition (Fig. 4i,k), of total loss after cluster-restricted silencing of essential inhibitory neurons (Fig. 4j,l), and of loss and Pearson correlation under targeted pathway blockade (Fig. 5p–s) used two-sided paired Student’s *t*-tests, with independently trained model instances (*n* = 5) providing the pairing factor. Across the three pairwise comparisons (S1 vs S2, S1 vs S3, S2 vs S3) within each panel, *p* values were adjusted by the Holm–Bonferroni procedure; exact corrected *p* values are reported numerically in the corresponding figure captions, and n.s. denotes *p* ≥ 0.05. Functional-connectivity matrices were compared by Pearson’s correlation across all neuropil pairs. The Pearson correlation between simulated and empirical neuropil activity reported in Figs. 3 and 5 was defined per neuropil and averaged across neuropils,

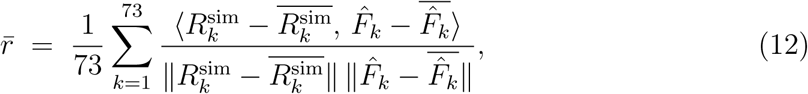

where (.)^-^ denotes the time average over the held-out test segment. Statistical tests were performed with scipy.stats^[66]^ and the powerlaw Python package^[65]^. The Pearson correlation was computed over the held-out test segment on the rate traces *R^sim^_k_* and *F*^^^*_k_* with each neuropil mean-centered before correlation; the same convention applied to every correlation value reported in the figures. With *n* = 5 paired model instances, the two-sided paired *t*-test with Holm–Bonferroni correction resolves standardized mean differences (Cohen’s *d_z_*) of approximately 2.5 at *α* = 0.05 and 80% power across the three cluster contrasts of each panel; effects smaller than this are not resolvable at the present sample size, and the reported *p* values in the 10*^−^*^3^–10*^−^*^5^ range reflect the large effect sizes evident in the underlying loss differences rather than fine statistical resolution.

## Data availability

The whole-brain calcium imaging recordings from *Drosophila* analyzed in this study are available from figshare at https://doi.org/10.6084/m9.figshare.13349282[9^]^. The preprocessed data and the trained model weights will be made available upon acceptance of the manuscript.

## Code availability

Code to reproduce the figures of this paper is built on our open-source BrainX ecosystem (https://github.com/chaobrain/brainx) and will be made available upon acceptance of the manuscript.

## Acknowledgements

This work was supported by the Young Scientists Fund of the National Natural Science Foundation of China (No. 3240070449, C.W.).

## Competing interests

The authors declare no competing interests.

## Notes

### Competing Interest Statement

The authors have declared no competing interest.

## References

1. Dorkenwald, S. et al. Neuronal wiring diagram of an adult brain. Nature 634, 124–138, DOI: 10.1038/s41586-024-07558-y (2024).

2. Shapson-Coe, A. et al. A petavoxel fragment of human cerebral cortex reconstructed at nanoscale resolution. Science 384, eadk4858, DOI: 10.1126/science.adk4858 (2024).

3. Consortium, M. Functional connectomics spanning multiple areas of mouse visual cortex. Nature 640, 435–447, DOI: 10.1038/s41586-025-08790-w (2025).

4. Urai, A. E., Doiron, B., Leifer, A. M. & Churchland, A. K. Large-scale neural recordings call for new insights to link brain and behavior. Nature neuroscience 25, 11–19, DOI: 10.1038/s41593-021-00980-9 (2022).

5. Trautmann, E. M. et al. Large-scale high-density brain-wide neural recording in nonhuman primates. Nature Neuroscience 28, 1562–1575, DOI: 10.1038/s41593-025-01976-5 (2025).

6. Findling, C. et al. Brain-wide representations of prior information in mouse decision-making. Nature 645, 192–200, DOI: 10.1038/s41586-025-09226-1 (2025).

7. Schlegel, P. et al. Whole-brain annotation and multi-connectome cell typing of Drosophila. Nature 634, 139–152, DOI: 10.1038/s41586-024-07686-5 (2024).

8. Mann, K., Gallen, C. L. & Clandinin, T. R. Whole-brain calcium imaging reveals an intrinsic functional network in Drosophila. Current Biology 27, 2389–2396.e4, DOI: 10.1016/j.cub.2017.06.076 (2017).

9. Turner, M. H., Mann, K. & Clandinin, T. R. The connectome predicts resting-state functional connectivity across the Drosophila brain. Current biology 31, 2386–2394.e3, DOI: 10.1016/j.cub.2021.03.004 (2021).

10. Fox, M. D. & Raichle, M. E. Spontaneous fluctuations in brain activity observed with functional magnetic resonance imaging. Nature reviews neuroscience 8, 700–711, DOI: 10.1038/nrn2201 (2007).

11. Smith, S. M. et al. Correspondence of the brain’s functional architecture during activation and rest. Proceedings national academy sciences 106, 13040–13045, DOI: 10.1073/pnas.0905267106 (2009).

12. Beggs, J. M. & Plenz, D. Neuronal avalanches in neocortical circuits. Journal neuroscience 23, 11167–11177, DOI: 10.1523/JNEUROSCI.23-35-11167.2003 (2003).

13. Shriki, O. et al. Neuronal avalanches in the resting MEG of the human brain. Journal Neuroscience 33, 7079–7090, DOI: 10.1523/JNEUROSCI.4286-12.2013 (2013).

14. Ponce-Alvarez, A., Jouary, A., Privat, M., Deco, G. & Sumbre, G. Whole-brain neuronal activity displays crackling noise dynamics. Neuron 100, 1446–1459.e6, DOI: 10.1016/j.neuron.2018.10.045 (2018).

15. Ito, K. et al. A systematic nomenclature for the insect brain. Neuron 81, 755–765, DOI: 10.1016/j.neuron.2013.12.017 (2014).

16. Buzsáki, G. & Mizuseki, K. The log-dynamic brain: how skewed distributions affect network operations. Nature reviews neuroscience 15, 264–278, DOI: 10.1038/nrn3687 (2014).

17. Gireesh, E. D. & Plenz, D. Neuronal avalanches organize as nested theta-and beta/gamma-oscillations during development of cortical layer 2/3. Proceedings National Academy Sciences 105, 7576–7581, DOI: 10.1073/pnas.0800537105 (2008).

18. Friedman, N. et al. Universal critical dynamics in high resolution neuronal avalanche data. Physical Review Letters 108, 208102, DOI: 10.1103/PhysRevLett.108.208102 (2012).

19. Sethna, J. P., Dahmen, K. A. & Myers, C. R. Crackling noise. Nature 410, 242–250, DOI: 10.1038/35065675 (2001).

20. Priesemann, V. et al. Spike avalanches in vivo suggest a driven, slightly subcritical brain state. Frontiers Systems Neuroscience 8, 108, DOI: 10.3389/fnsys.2014.00108 (2014).

21. Levina, A. & Priesemann, V. Subsampling scaling. Nature Communications 8, 15140, DOI: 10.1038/ncomms15140 (2017).

22. van den Heuvel, M. P. & Sporns, O. Rich-club organization of the human connectome. Journal Neuroscience 31, 15775–15786, DOI: 10.1523/JNEUROSCI.3539-11.2011 (2011).

23. van den Heuvel, M. P. & Sporns, O. Network hubs in the human brain. Trends Cognitive Sciences 17, 683–696, DOI: 10.1016/j.tics.2013.09.012 (2013).

24. Bassett, D. S. & Sporns, O. Network neuroscience. Nature Neuroscience 20, 353–364, DOI: 10.1038/nn.4502 (2017).

25. Lin, A. et al. Network statistics of the whole-brain connectome of Drosophila. Nature 634, 153–165, DOI: 10.1038/s41586-024-07968-y (2024).

26. Aimon, S. et al. Fast near-whole–brain imaging in adult Drosophila during responses to stimuli and behavior. PLoS biology 17, e2006732, DOI: 10.1371/journal.pbio.2006732 (2019).

27. Tang, S. & Juusola, M. Intrinsic activity in the fly brain gates visual information during behavioral choices. PLOS ONE 5, 1–19, DOI: 10.1371/journal.pone.0014455 (2011).

28. Akin, O., Bajar, B. T., Keles, M. F., Frye, M. A. & Zipursky, S. L. Cell-type-specific patterned stimulus-independent neuronal activity in the Drosophila visual system during synapse formation. Neuron 101, 894–904.e5, DOI: 10.1016/j.neuron.2019.01.008 (2019).

29. Donlea, J. M., Pimentel, D. & Miesenböck, G. Neuronal machinery of sleep homeostasis in Drosophila. Neuron 81, 860–872, DOI: 10.1016/j.neuron.2013.12.013 (2014).

30. Tainton-Heap, L. A. L. et al. A paradoxical kind of sleep in Drosophila melanogaster. Current Biology 31, 578–590.e6, DOI: 10.1016/j.cub.2020.10.081 (2021).

31. van Swinderen, B. & Greenspan, R. J. Salience modulates 20–30 Hz brain activity in Drosophila. Nature Neuroscience 6, 579–586, DOI: 10.1038/nn1054 (2003).

32. Behnia, R., Clark, D. A., Carter, A. G., Clandinin, T. R. & Desplan, C. Processing properties of ON and OFF pathways for Drosophila motion detection. Nature 512, 427–430, DOI: 10.1038/nature13427 (2014).

33. Strother, J. A. et al. The emergence of directional selectivity in the visual motion pathway of Drosophila. Neuron 94, 168–182.e10, DOI: 10.1016/j.neuron.2017.03.010 (2017).

34. Takemura, S.-y., et al. A visual motion detection circuit suggested by Drosophila connectomics. Nature 500, 175–181, DOI: 10.1038/nature12450 (2013).

35. Nern, A., Pfeiffer, B. D. & Rubin, G. M. Optimized tools for multicolor stochastic labeling reveal diverse stereotyped cell arrangements in the fly visual system. Proceedings National Academy Sciences 112, E2967–E2976, DOI: 10.1073/pnas.1506763112 (2015).

36. Maisak, M. S. et al. A directional tuning map of Drosophila elementary motion detectors. Nature 500, 212–216, DOI: 10.1038/nature12320 (2013).

37. Wolff, T. & Rubin, G. M. Neuroarchitecture of the Drosophila central complex: A catalog of nodulus and asymmetrical body neurons and a revision of the protocerebral bridge catalog. Journal Comparative Neurology 526, 2585–2611, DOI: 10.1002/cne.24512 (2018).

38. Turner-Evans, D. B. et al. The neuroanatomical ultrastructure and function of a biological ring attractor. Neuron 108, 145–163.e10, DOI: 10.1016/j.neuron.2020.08.006 (2020).

39. Aso, Y. et al. The neuronal architecture of the mushroom body provides a logic for associative learning. elife 3, e04577, DOI: 10.7554/eLife.04577 (2014).

40. Mauss, A. S. et al. Neural circuit to integrate opposing motions in the visual field. Cell 162, 351–362, DOI: 10.1016/j.cell.2015.06.035 (2015).

41. Chou, Y.-H. et al. Diversity and wiring variability of olfactory local interneurons in the *Drosophila* antennal lobe. Nature Neuroscience 13, 439–449, DOI: 10.1038/nn.2489 (2010).

42. Takemura, S.-y., et al. The comprehensive connectome of a neural substrate for ‘ON’ motion detection in Drosophila. eLife 6, e24394, DOI: 10.7554/eLife.24394 (2017).

43. Molina-Obando, S. et al. ON selectivity in the Drosophila visual system is a multi-synaptic process involving both glutamatergic and GABAergic inhibition. eLife 8, e49373, DOI: 10.7554/eLife.49373 (2019).

44. Shiu, P. K. et al. A Drosophila computational brain model reveals sensorimotor processing. Nature 634, 210–219, DOI: 10.1038/s41586-024-07763-9 (2024).

45. Lappalainen, J. K. et al. Connectome-constrained networks predict neural activity across the fly visual system. Nature 634, 1132–1140, DOI: 10.1038/s41586-024-07939-3 (2024).

46. Deco, G., Kringelbach, M. L., Jirsa, V. K. & Ritter, P. The dynamics of resting fluctuations in the brain: metastability and its dynamical cortical core. Scientific reports 7, 3095, DOI: 10.1038/s41598-017-03073-5 (2017).

47. Lueckmann, J.-M. et al. Zapbench: A benchmark for whole-brain activity prediction in zebrafish. In International Conference on Learning Representations, vol. 2025, 41450–41487 (2025).

48. Pereda, A. E. Electrical synapses and their functional interactions with chemical synapses. Nature Reviews Neuroscience 15, 250–263, DOI: 10.1038/nrn3708 (2014).

49. de Ruyter van Steveninck, R. R. & Laughlin, S. B. The rate of information transfer at graded-potential synapses. Nature 379, 642–645, DOI: 10.1038/379642a0 (1996).

50. Bargmann, C. I. Beyond the connectome: how neuromodulators shape neural circuits. Bioessays 34, 458–465, DOI: 10.1002/bies.201100185 (2012).

51. Fan, J. et al. Prominent involvement of acetylcholine dynamics in stable olfactory representation across the Drosophila brain. Nature Communications 16, 8638, DOI: 10.1038/s41467-025-63823-2 (2025).

52. Vogelstein, J. T. et al. Fast nonnegative deconvolution for spike train inference from population calcium imaging. Journal Neurophysiology 104, 3691–3704, DOI: 10.1152/jn.01073.2009 (2010).

53. Pnevmatikakis, E. A. et al. Simultaneous denoising, deconvolution, and demixing of calcium imaging data. Neuron 89, 285–299, DOI: 10.1016/j.neuron.2015.11.037 (2016).

54. Chen, T.-W. et al. Ultrasensitive fluorescent proteins for imaging neuronal activity. Nature 499, 295–300, DOI: 10.1038/nature12354 (2013).

55. Hansen, P. C. Analysis of discrete ill-posed problems by means of the L-curve. SIAM Review 34, 561–580, DOI: 10.1137/1034115 (1992).

56. Beck, A. & Teboulle, M. A fast iterative shrinkage-thresholding algorithm for linear inverse problems. SIAM Journal on Imaging Sciences 2, 183–202, DOI: 10.1137/080716542 (2009).

57. Werbos, P. J. Backpropagation through time: What it does and how to do it. Proc. IEEE 78, 1550–1560, DOI: 10.1109/5.58337 (1990).

58. Wang, C. et al. Model-agnostic linear-memory online learning in spiking neural networks. Nature Communications DOI: 10.1038/s41467-026-68453-w (2026).

59. Kingma, D. P. & Ba, J. Adam: a method for stochastic optimization. In Proceedings of the 3rd International Conference on Learning Representations (ICLR) (2015).

60. Wang, C. et al. Brainpy, a flexible, integrative, efficient, and extensible framework for general-purpose brain dynamics programming. eLife 12, DOI: 10.7554/elife.86365 (2023).

61. Wang, C., et al. A differentiable brain simulator bridging brain simulation and brain-inspired computing. In The Twelfth International Conference on Learning Representations (2024).

62. Wang, C., He, S., Luo, S., Huan, Y. & Wu, S. Integrating physical units into high-performance AI-driven scientific computing. Nature Communications 16, DOI: 10.1038/s41467-025-58626-4 (2025).

63. Romano, S. A. et al. An integrated calcium imaging processing toolbox for the analysis of neuronal population dynamics. PLoS Computational Biology 13, e1005526, DOI: 10.1371/journal.pcbi.1005526 (2017).

64. Clauset, A., Shalizi, C. R. & Newman, M. E. J. Power-law distributions in empirical data. SIAM Review 51, 661–703, DOI: 10.1137/070710111 (2009).

65. Alstott, J., Bullmore, E. & Plenz, D. powerlaw: a Python package for analysis of heavy-tailed distributions. PLoS ONE 9, e85777, DOI: 10.1371/journal.pone.0085777 (2014).

66. Virtanen, P. et al. SciPy 1.0: fundamental algorithms for scientific computing in Python. Nature Methods 17, 261–272, DOI: 10.1038/s41592-019-0686-2 (2020).

