## Supplemental Information for "Connectome-constrained modeling identifies neurons and synapses that sustain spontaneous activity in *Drosophila*"

### 1 Supplementary Figures

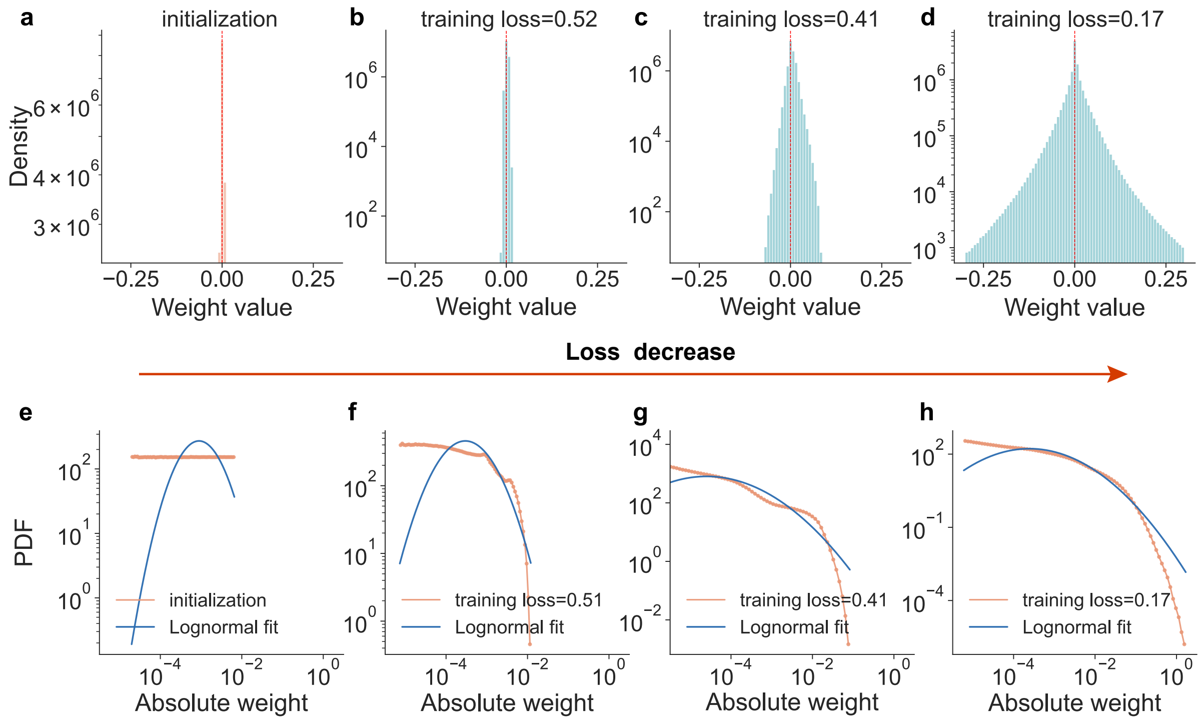


**Supplementary Figure 1. The effective synaptic-weight distribution broadens during training and converges to a lognormal shape as training loss decreases.** Effective synaptic weights at four representative training stages (initialization and training-loss snapshots ∼0.52, 0.41, and 0.17), pooled across $n = 5$ independently trained models ($\sim 1.5 \times 10^7$ synapses each). The orange "Loss decrease" arrow between the two rows indicates progression along the training trajectory. **a–d,** Histograms of signed effective synaptic weights at each training stage (linear weight axis; log-scale density in **b–d**, linear in **a**). Red dashed line marks zero. The initially tightly peaked distribution at zero (**a**) progressively broadens into a symmetric, heavy-tailed shape spanning excitatory (positive) and inhibitory (negative) values across several orders of magnitude (**b–d**). **e–h,** Probability density function (PDF) of absolute synaptic weights at the same four stages on log–log axes (orange), overlaid with the maximum-likelihood truncated lognormal fit (blue; Clauset et al. framework[1], `powerlaw` package[2]). The empirical distribution departs strongly from the lognormal reference at initialization (**e**) and progressively converges toward it as training loss decreases (**f–g**), with the converged distribution (**h**) tracking the lognormal fit across roughly four orders of magnitude in weight. Panel **h** corresponds to the converged distribution quantified in Fig. 1h.

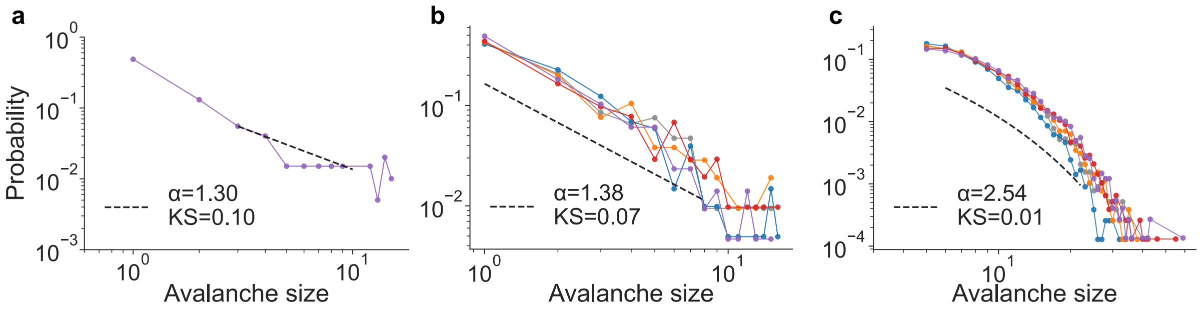


**Supplementary Figure 2. Avalanche-size distributions are power-law distributed in both empirical and simulated resting-state activity, paralleling the avalanche-duration statistics in Fig. 1i–k.** Probability distributions of avalanche sizes (number of supra-threshold events per avalanche) on log–log axes, with the maximum-likelihood truncated power-law fit shown as a dashed black line and the corresponding exponent $\alpha$ and Kolmogorov–Smirnov goodness-of-fit statistic ($KS$) annotated in each panel (Clauset et al. framework[1], `powerlaw` package[2]; see Supplementary Note 2). **a,** Avalanche-size distribution from empirical neuropil-level calcium imaging. Power-law exponent $\alpha = 1.30$, $KS = 0.10$ ($p = 1.3 \times 10^{-3}$ vs. lognormal; Supplementary Note 3). **b,** Avalanche-size distribution from simulated neuropil activity ($n = 5$ independently trained models, one curve per model). Power-law exponent $\alpha = 1.38$, $KS = 0.07$ ($p = 4.15 \times 10^{-12}$ vs. lognormal; Supplementary Note 3), closely matching the empirical neuropil-level statistic in **a**. **c,** Single-neuron avalanche-size distribution from simulated activity ($n = 5$ independently trained models, one curve per model). Power-law exponent $\alpha = 2.54$, $KS = 0.01$ ($p = 6.13 \times 10^{-13}$ vs. lognormal; Supplementary Note 3).

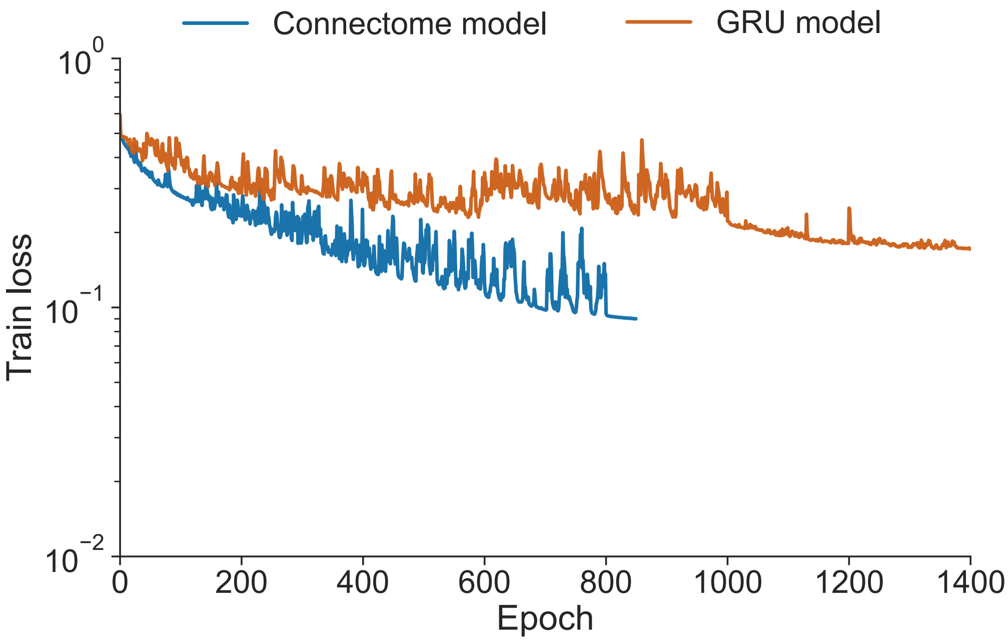


**Supplementary Figure 3. The connectome-constrained model and the parameter-matched GRU both converge under the same next-step training loss.** Training loss as a function of optimization epoch for the connectome-constrained model (blue) and the parameter-matched GRU baseline (orange), both trained on neuropil activity under the next-step mean-squared-error loss (Eq. (7); see Methods). The $y$-axis is logarithmic. Both models reach a low and stable training loss, so the divergent rollout behavior in Fig. 2a,b reflects a difference in generalization beyond the training window, not a failure of the GRU to fit the training segment.

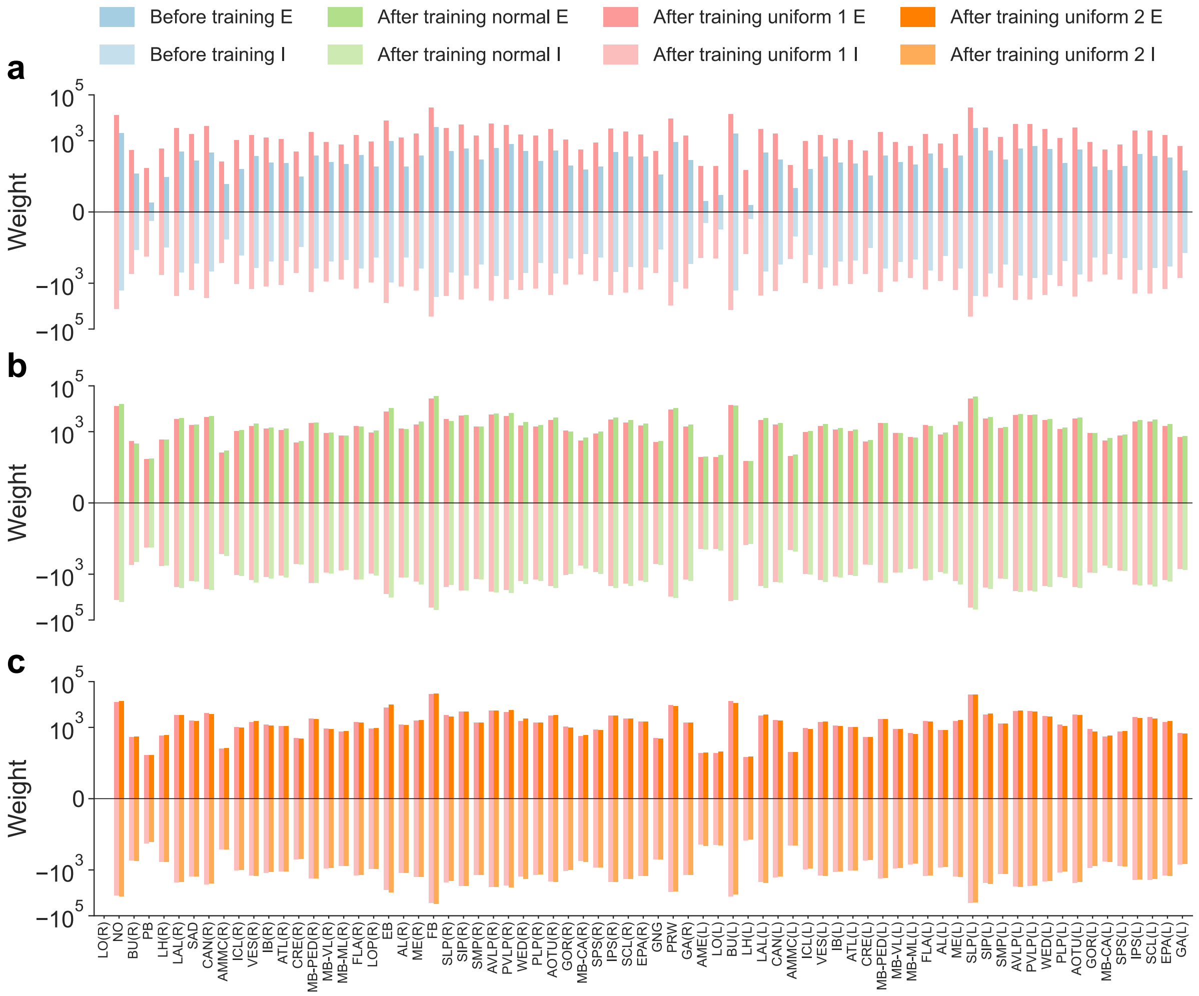


**Supplementary Figure 4. Learned neuropil-level weight distributions are robust to initialization and reproducible across independent random seeds. a,** Distributions of $\log_{10}$(total post-synaptic weight) per neuropil for excitatory (top) and inhibitory (bottom) populations, comparing the initial values ("Initial") with the converged values after training under a uniform prior ("Uniform 1"). Training drives both populations substantially away from their initial states. **b,** Per-neuropil percentage difference between converged weights obtained under a truncated-normal prior ("Normal") and those obtained under Uniform 1, for excitatory (top) and inhibitory (bottom) populations. Converged weights remain within $\pm 5\%$ across prior families, indicating that the learned weights are largely independent of the initialization family. **c,** Same as **b**, but comparing converged weights obtained under a second uniform prior with different random seeds ("Uniform 2") against those obtained under Uniform 1. Converged weights remain within $\pm 5\%$ across seeds.

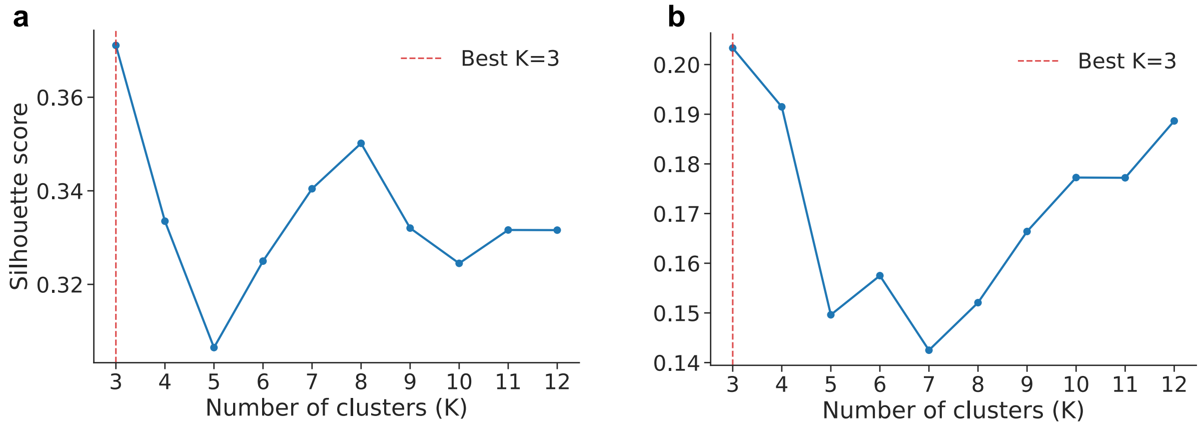


**Supplementary Figure 5. Silhouette-coefficient curves motivate the $k = 3$ source- and target-cluster solutions.** Silhouette coefficient as a function of cluster number $k$ for hierarchical agglomerative clustering (Euclidean distance, average linkage) of the directed effectome matrix (Fig. 3b), evaluated over $k \in \{2, \ldots, 12\}$. **a,** Source clustering: silhouette score for partitions of the row vectors of the effectome matrix. The maximum at $k = 3$ (red dashed line) defines the three-cluster source partition (S1, S2, S3) used throughout the main text. **b,** Target clustering: silhouette score for partitions of the column vectors of the effectome matrix. The maximum at $k = 3$ defines the three-cluster target partition (T1, T2, T3). In both partitions, $k = 3$ outperforms all alternatives in $\{2, 4, 5, \ldots, 12\}$, supporting the choice used in Fig. 3 and downstream analyses.

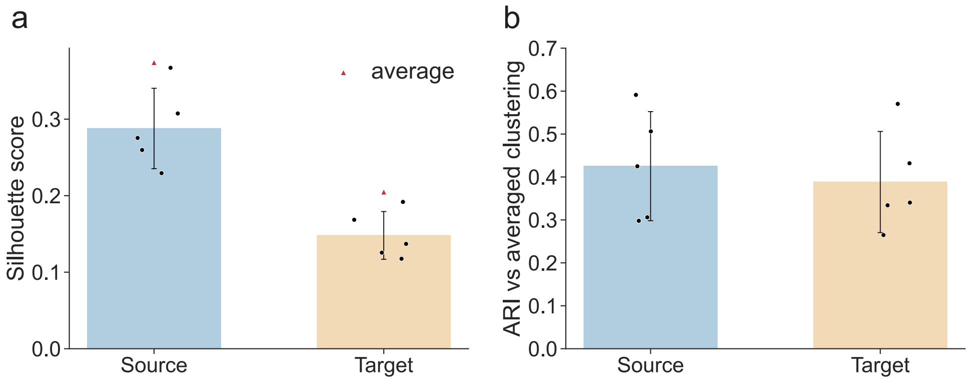


**Supplementary Figure 6. The $k = 3$ source and target partitions are reproducible across independently trained models.** For each of the $n = 5$ independently trained models, the directed effectome matrix (Fig. 3b) was clustered with the same hierarchical agglomerative procedure used for the averaged matrix (Euclidean distance, average linkage, $k = 3$; Methods), and the resulting per-model partitions were compared with the partition obtained on the averaged effectome. **a,** Silhouette coefficient at $k = 3$ for the source partition (blue) and the target partition (orange). Black dots, per-model values ($n = 5$); bars, mean; error bars, standard deviation; red triangles, silhouette coefficient of the clustering computed on the averaged effectome. The averaged clustering matches or exceeds the per-model mean for both partitions, so averaging across models does not blur cluster structure. **b,** Adjusted Rand Index (ARI) between each per-model clustering and the averaged clustering, for the source (blue) and target (orange) partitions. Both partitions lie well above the chance level (ARI $\approx 0$) expected for unrelated $k = 3$ assignments, indicating substantial agreement between per-model and averaged clusterings and supporting the use of a single averaged clustering throughout the main text.

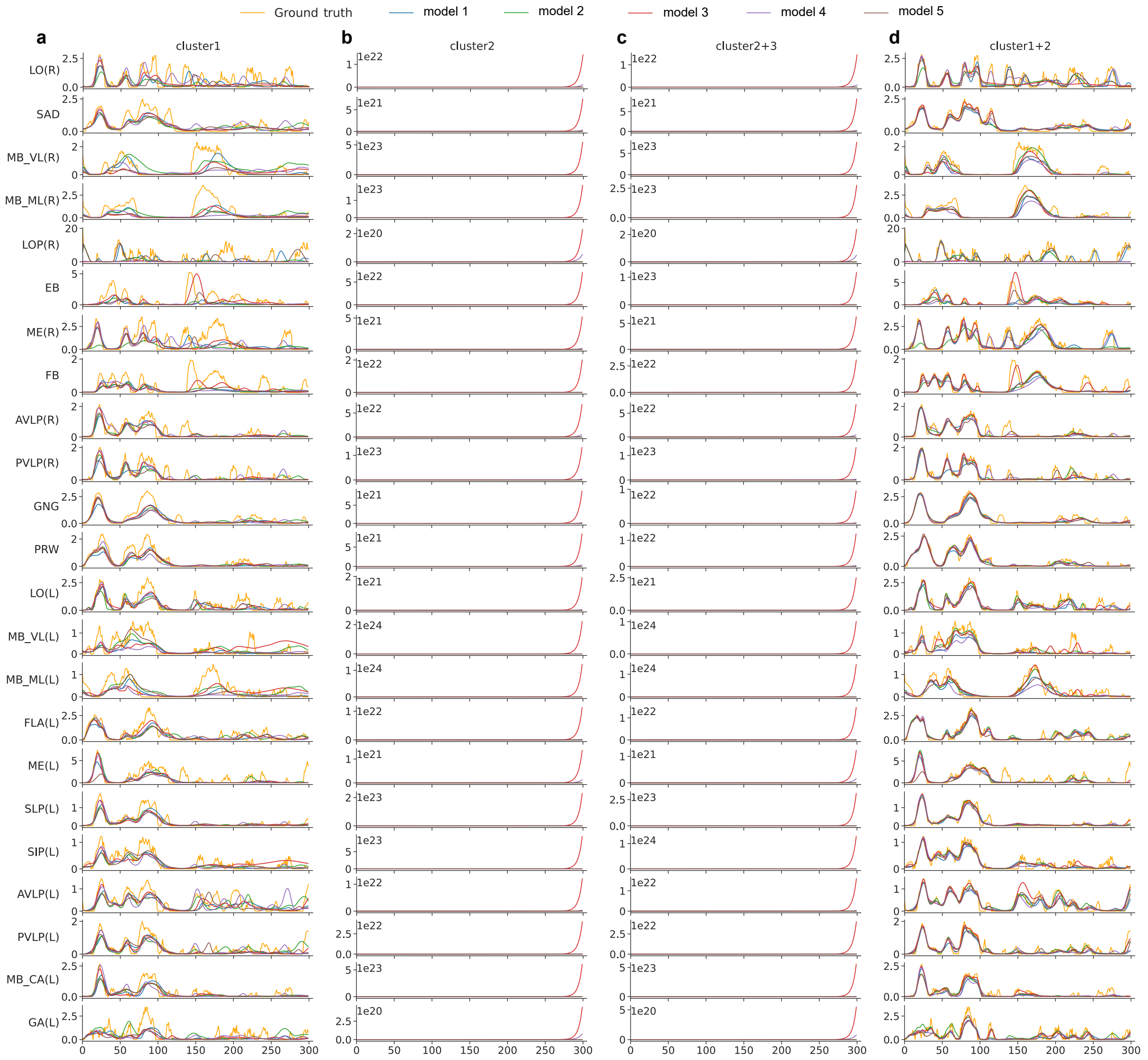


**Supplementary Figure 7. Whole-brain simulated traces under selective source-cluster preservation.** Simulated activity (colored) overlaid on empirical ground truth (gray) for the first 25 neuropils under the four cluster-preservation conditions of Fig. 3d–g, extending the four-neuropil summary in the main figure to the full brain. **a,** S1 alone preserved (other clusters ablated *in silico*; cf. Fig. 3d). **b,** S2 alone preserved (cf. Fig. 3e). **c,** S2+S3 preserved, S1 ablated (cf. Fig. 3f). **d,** S1+S2 preserved, S3 ablated (cf. Fig. 3g). Each subpanel shows one neuropil; rows labeled at left follow the Ito reference atlas[3] (see Supplementary Table 2). Vertical dashed line marks the boundary between the training set (left) and the held-out test set (right). Traces show one representative simulation per condition from a single trained model; $n = 5$ independently trained models gave consistent results.

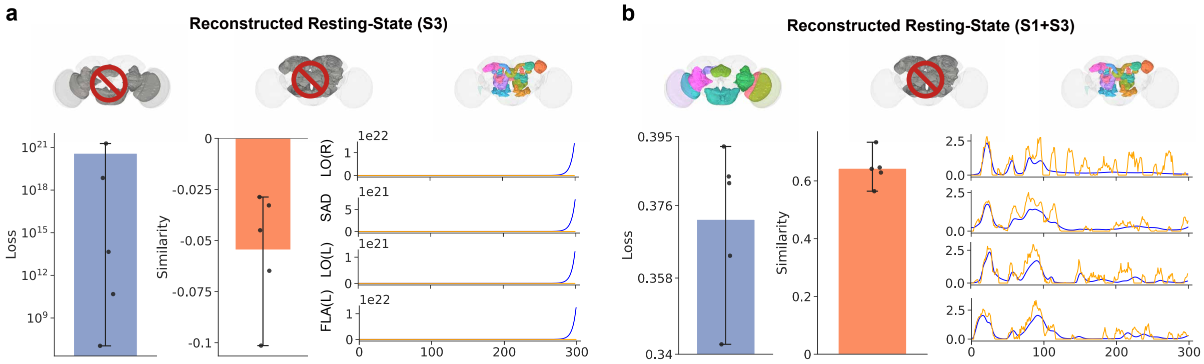


**Supplementary Figure 8. Additional source-cluster preservation conditions confirm that S1 is necessary and that S2 and S3 contribute interchangeable complementary detail.** Each panel reports test-set prediction loss summed over all neuropils (left, blue bar), Pearson correlation between simulated and empirical neuropil activity (middle, orange bar), and representative simulated traces for four example neuropils (LO(R), SAD, LO(L), FLA(L); predicted in blue, empirical ground truth in orange). Brain icons indicate intact and ablated clusters (red prohibition symbol). Bars, mean ± std.; $n = 5$ independently trained models (one independent simulation per model per condition). **a,** S3 alone preserved (S1 and S2 ablated *in silico*). The rate dynamics fail to remain bounded under this perturbation (rates diverge to $\sim 10^{21}$ in the loss metric; Pearson $r \sim -0.05$), mirroring the instability observed when only S2 is preserved (cf. Fig. 3e); preserving any non-S1 cluster alone is therefore insufficient to sustain resting-state activity. **b,** S1+S3 preserved (S2 ablated). Stable resting-state activity is restored (loss ∼0.38; Pearson $r \sim 0.64$), comparable to the S1+S2 condition (cf. Fig. 3g), indicating that S2 and S3 contribute interchangeable complementary detail once the S1 core is preserved. Neuropil abbreviations follow the Ito reference atlas[3]; see Supplementary Table 2.

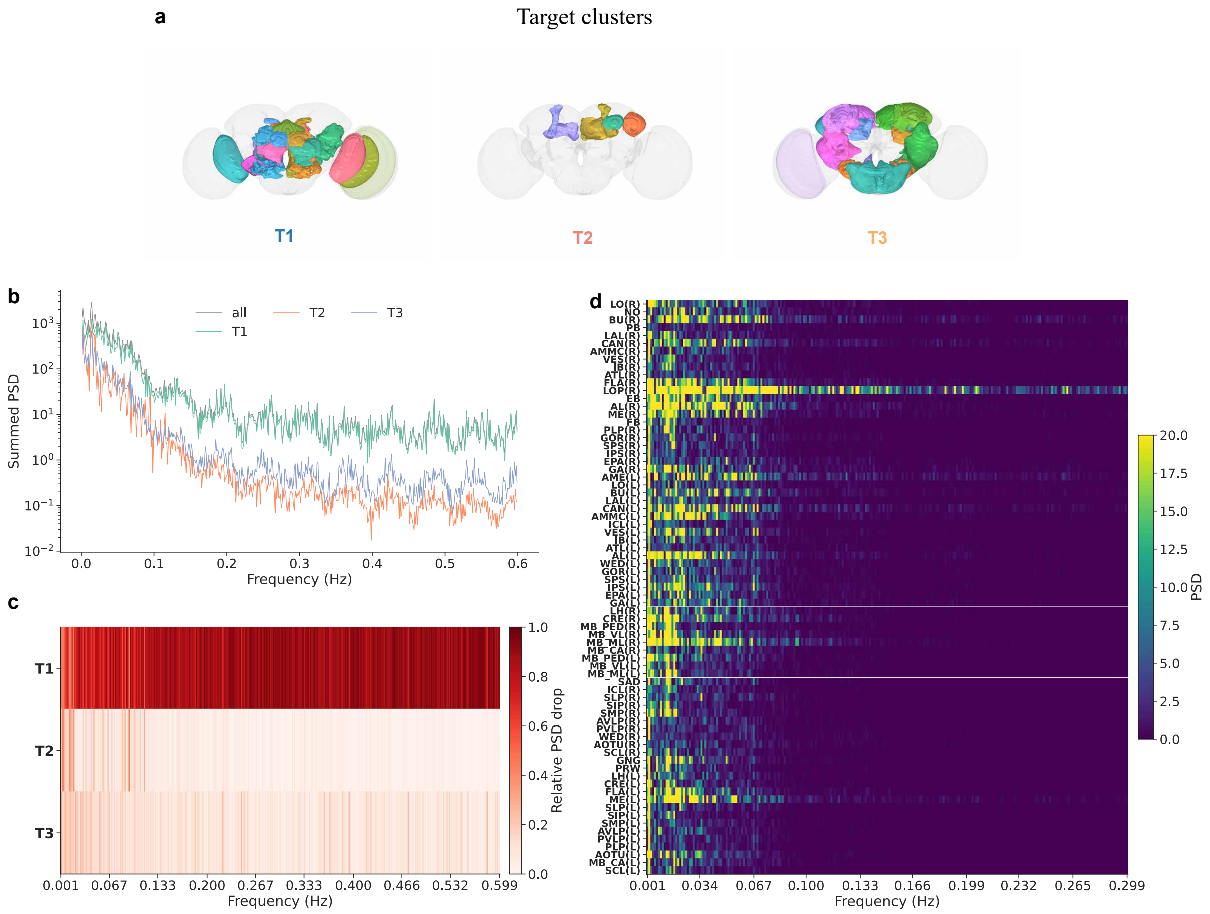


**Supplementary Figure 9. Spectral characterization of target-neuropil clusters in the trained connectome-constrained model.** **a,** Anatomical layout of the three target-neuropil clusters T1, T2, and T3 identified by clustering the columns of the directed effectome matrix (Fig. 3b), color-coded and projected onto the standard *Drosophila* brain atlas. **b,** Cluster-summed power spectral density (PSD) of simulated neuropil activity from the trained model. Traces, total PSD summed across all neuropils (*all*, gray) and separately within target clusters T1 (green), T2 (orange), and T3 (purple), as a function of frequency (0–0.6 Hz, log-scaled $y$-axis). $n = 5$ independently trained models. **c,** Heatmap of the relative PSD drop induced by *in silico* ablation of each target cluster, resolved across frequency bins from 0.001 to 0.599 Hz. Rows, ablated clusters (T1, T2, T3); color, fractional reduction in summed PSD relative to the intact model (0–1, white to dark red). $n = 5$ trained models. **d,** Per-neuropil PSD heatmap of simulated activity. Rows, individual neuropils (labels at left); columns, frequency bins from 0.001 to 0.299 Hz; color, PSD magnitude (0–20). Horizontal white separators group neuropils by target-cluster assignment. The low-frequency dominance in **b,d** partly reflects the GCaMP6s calcium kernel applied to the training target, which acts as a ~10-second low-pass filter ($\tau_{\mathrm{Ca}} = 1.5\,\mathrm{s}$; Methods, Eq. (1)). The analogy with vertebrate resting-state infraslow fluctuations[4,5] should be read with this confound in mind. Neuropil abbreviations follow the Ito reference atlas[3]; see Supplementary Table 2.

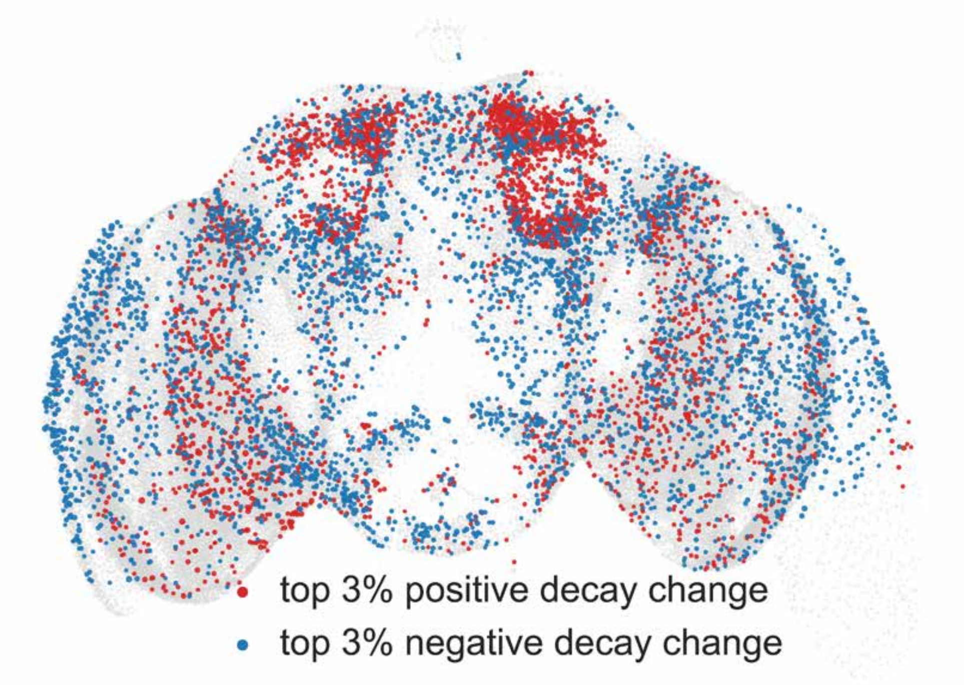


**Supplementary Figure 10. The $\Delta\tau$-decrease and $\Delta\tau$-increase tail subpopulations occupy spatially compact, hemisphere-biased regions of the standard *Drosophila* brain.** Somata of the two tail subpopulations identified in Fig. 4a, projected onto the standard *Drosophila* brain template. Red, top-3% positive $\Delta\tau$ tail (lengthened-timescale neurons, "increase tail"); blue, top-3% negative $\Delta\tau$ tail (shortened-timescale neurons, "decrease tail"). Soma coordinates of each FlyWire neuron were registered to the template and pooled across $n = 5$ independently trained models. Neither subpopulation distributes uniformly; both concentrate in spatially compact, hemisphere-biased ensembles, consistent with the anatomically selective remodeling of intrinsic timescales reported in Fig. 4 and quantified at the cell-type level in Fig. 6a.

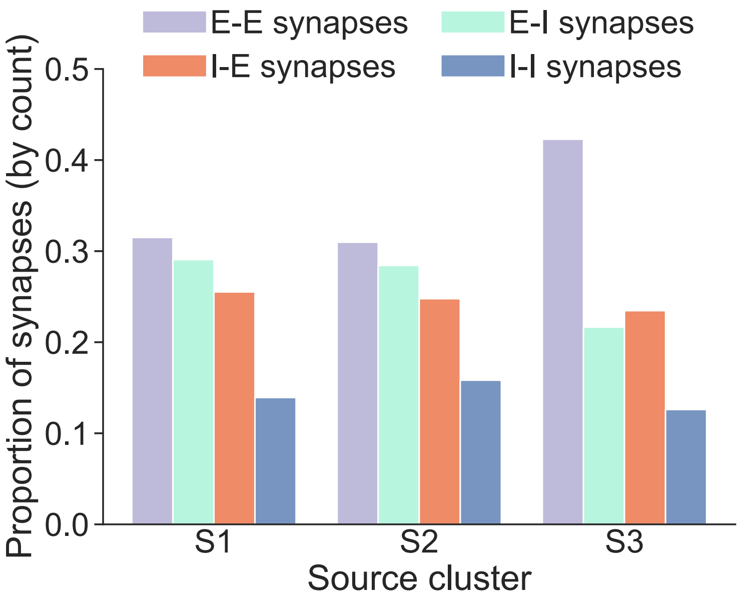


**Supplementary Figure 11. Within-cluster pathway-class composition is uniform across source clusters.** For each source cluster (S1, S2, S3), bars give the within-cluster fraction of intra-cluster synapses falling into each of the four polarity classes (E–E, E–I, I–E, I–I); the four bars sum to one within each cluster. The relative mix is nearly identical in S1, S2, and S3, complementing the cross-cluster view in Fig. 5b: no pathway class is preferentially enriched in any one cluster, and pathway-class composition therefore does not distinguish S1 from S2 or S3.

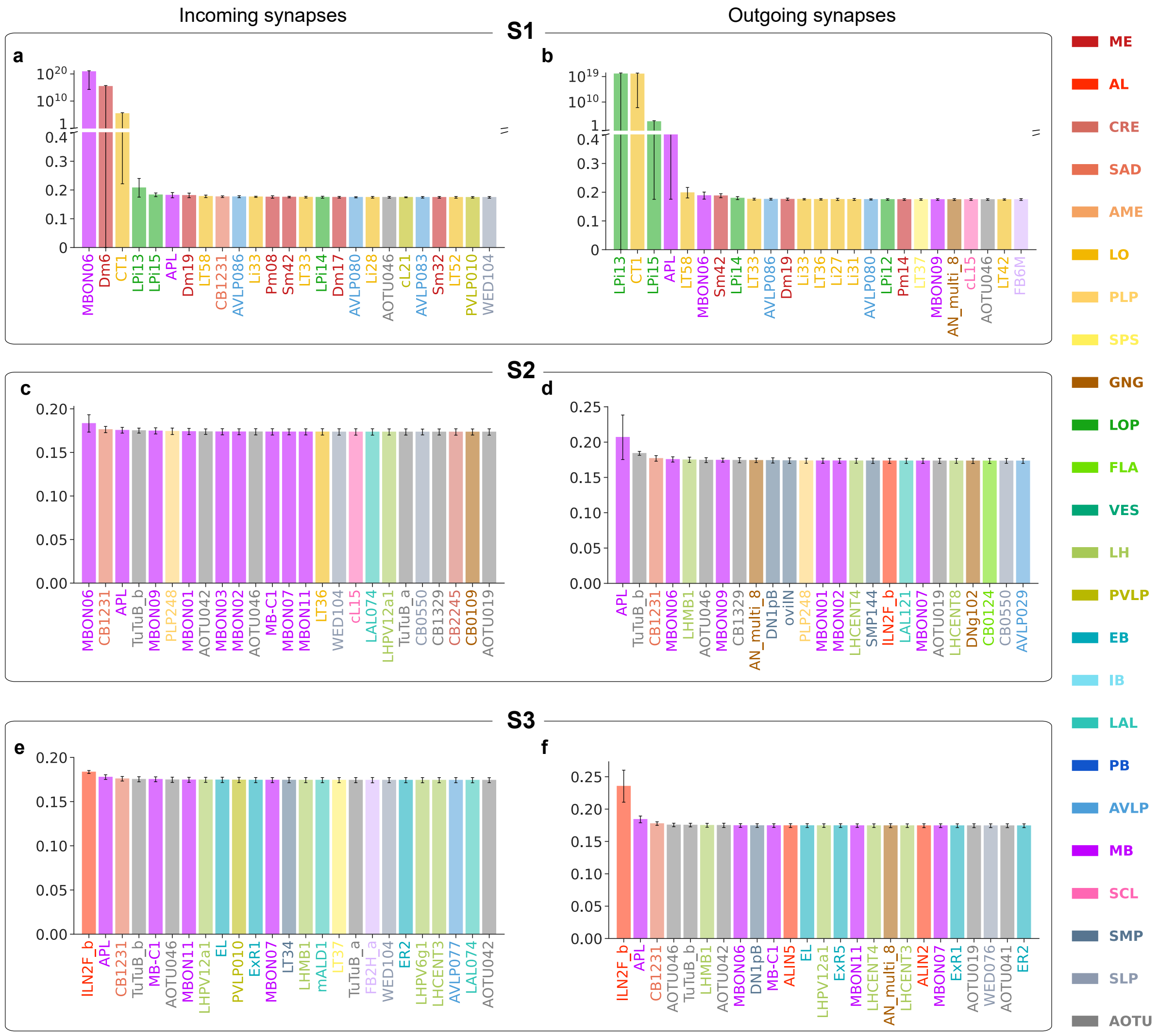


**Supplementary Figure 12. Per-cell-type synaptic-block loss for the input and output sides of the top-5% weighted inhibitory hub, resolved by source cluster (S1, S2, S3). a,** Input-side per-cell-type synaptic-block loss for the S1 cluster. For each presynaptic partner cell type sending S1-routed synapses onto the top-5% weighted inhibitory neurons of Fig. 4g–l, the synapses it contributes to the hub are set to zero and the model is re-run; the bar gives the resulting prediction loss against empirical activity (Fig. 5p,r protocol). Top loss values are shown on a broken axis (inset to log scale). **b,** Output-side per-cell-type synaptic-block loss for the S1 cluster, constructed as in **a** but for output synapses from the hub to each postsynaptic target cell type. **c,d,** As in **a,b** but restricted to synapses routed through the S2 cluster: input-side block loss (**c**) and output-side block loss (**d**). **e,f,** As in **a,b** but restricted to synapses routed through the S3 cluster: input-side block loss (**e**) and output-side block loss (**f**). Only the 30 most weight-contributing cell types per panel are shown. Bars, mean across $n = 5$ trained models; whiskers, std. Cell-type labels are color-coded by primary neuropil of innervation following Fig. 6 and the Ito atlas[3]; cell-type names follow the FlyWire annotation[6]. Block-loss axes share a common scale across S1, S2, and S3 to allow direct comparison. The figure resolves the reciprocal E–I/I–E motif of Figs. 3 and 5 at the cell-type level and shows that the motif is carried by S1: the synapses whose removal collapses the loop are concentrated on a small set led by MBON06 on the input side (**a**) and the LPi opponent-inhibition cells on the output side (**b**); equivalent cell-type-resolved block within S2 (**c,d**) or S3 (**e,f**) does not raise the loss above its baseline floor for any cell type.

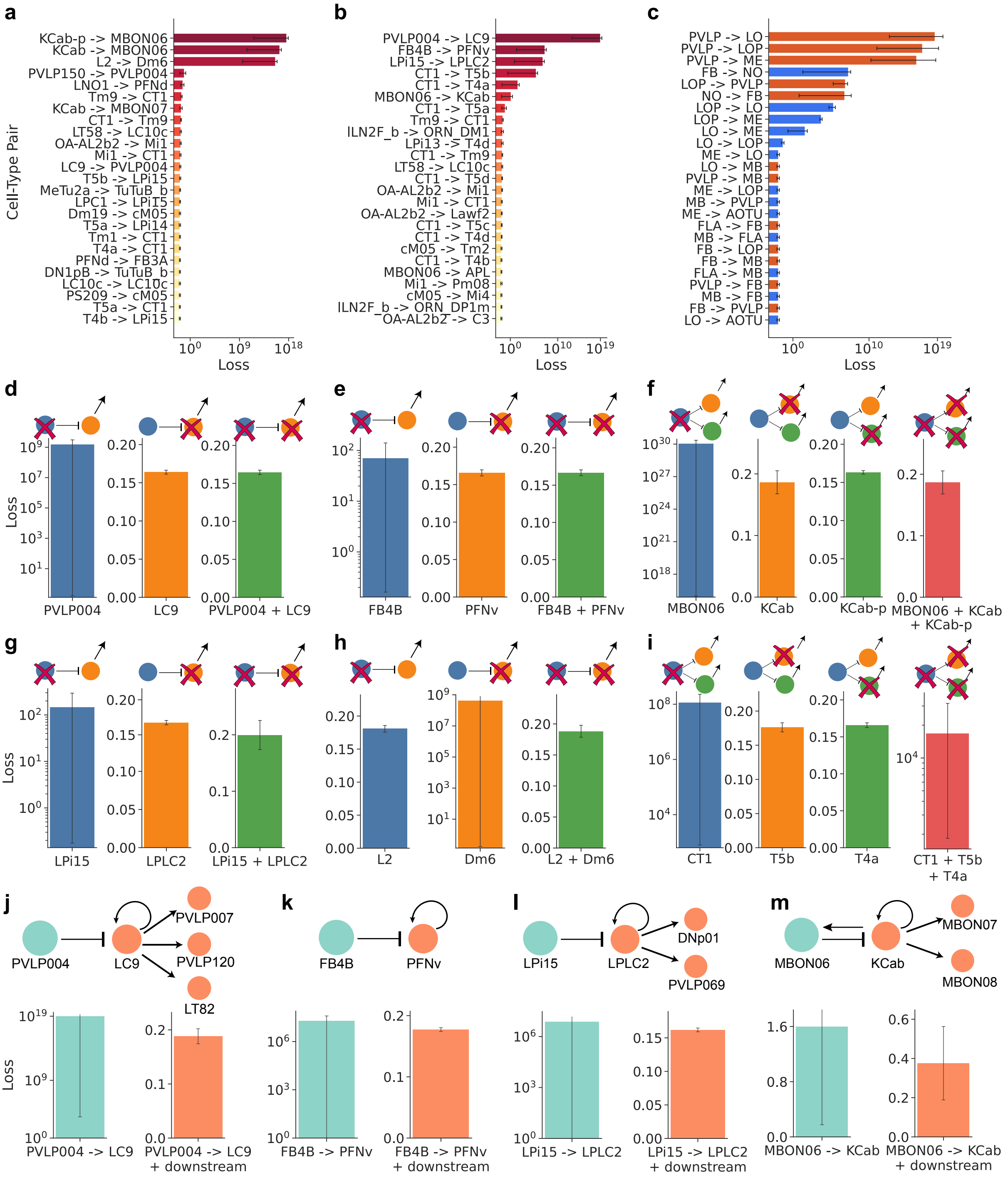


**Supplementary Figure 13. Single- versus combined-silencing of cell types and synaptic projections resolves the essential inhibitory hubs into reciprocal excitatory–inhibitory partnerships. a,** Input-side per-pair silencing loss. For each presynaptic cell type that projects onto an essential cell type of Fig. 6f, the synapses of that ordered cell-type pair were set to zero and the model was re-run on the held-out test segment; the bar gives the resulting prediction loss. Pairs are ranked by loss (top entries KCab-p → MBON06, KCab → MBON06, L2 → Dm6, PVLP150 → PVLP004). **b,** Output-side per-pair silencing loss, as in **a** but for synapses from each essential cell type onto its postsynaptic partners (top entries PVLP004 → LC9, FB4B → PFNv, LPi15 → LPLC2, CT1 → T5b, MBON06 → KCab).

**Supplementary Figure 13.** **c,** Neuropil-pair silencing loss. Synapses were grouped by the directed neuropil pair (source $\rightarrow$ target) of their endpoints together with their polarity, then silenced as a group; bars give the summed loss, colored by the polarity of the silenced synapses (red, excitatory; blue, inhibitory). **d–i,** Single- versus combined-cell-type silencing. In each panel the inhibitory hub cell type, its excitatory partner cell type(s), and their combination are silenced in turn (clamped to zero rate). Silencing the inhibitory hub alone drives the loss to unbounded values (log axes); silencing the excitatory partner alone leaves the loss near baseline ($\sim$0.16–0.2); silencing the hub together with its partner(s) returns the loss to baseline. **d,** PVLP004 with LC9. **e,** FB4B with PFNv. **f,** MBON06 with KCab and KCab-p. **g,** LPi15 with LPLC2. **h,** Dm6 with L2. **i,** CT1 with T5b and T4a; here the combination lowers the loss by several orders of magnitude but not fully to baseline. **j–m,** Single- versus combined-projection silencing, as in **d–i** but blocking synaptic projections rather than whole cells. Blocking the inhibitory hub-to-partner projection alone (left) drives the loss to unbounded values; additionally blocking the partner's downstream excitatory projections (right; schematic above each panel) returns the loss to baseline. **j,** PVLP004 $\dashv$ LC9, with LC9 $\rightarrow$ $\{$PVLP007, PVLP120, LT82$\}$. **k,** FB4B $\dashv$ PFNv, with the downstream projections of PFNv. **l,** LPi15 $\dashv$ LPLC2, with LPLC2 $\rightarrow$ $\{$DNp01, PVLP069$\}$. **m,** MBON06 $\dashv$ KCab, with KCab $\rightarrow$ $\{$MBON07, MBON08$\}$. Bars, mean $\pm$ std.; $n = 5$ independently trained models. Cell-type names follow the FlyWire annotation[6]; neuropil abbreviations follow the Ito atlas[3]. Across hubs, the indispensability of each essential inhibitory cell type tracks a reciprocal partnership with a specific excitatory cell type: removing the inhibition releases the excitatory partner, and removing that partner, whether as a cell (**d–i**) or as a projection (**j–m**), reabsorbs the runaway.

#### 2 Supplementary Tables

**Supplementary Table 1. Neuropil cluster membership.** Source clusters (S1–S3) were obtained by hierarchical agglomerative clustering of the rows of the directed effectome matrix (Fig. 3b); target clusters (T1–T3) by clustering of its columns. Neuropil abbreviations follow the Ito reference atlas[3]; (L), (R), and (L/R) denote left, right, and bilateral, respectively.

| Cluster | $n$ | Member neuropils |
|---|---|---|
| S1 | 14 | ME(L/R), LO(L/R), LOP(R), AVLP(L/R), PVLP(L/R), PLP(L), GNG, PRW, FB, MB-ML(L) |
| S2 | 33 | SMP(L/R), SAD, SPS(L/R), SIP(L/R), IPS(L/R), LAL(L/R), VES(L/R), CRE(L/R), ICL(L/R), AOTU(R), FLA(L/R), PLP(R), SLP(L/R), SCL(L/R), IB(L/R), WED(L/R), LH(L), MB-ML(R), MB-VL(L/R) |
| S3 | 26 | AL(L/R), LH(R), GOR(L/R), NO, EB, PB, AOTU(L), EPA(L/R), AMMC(L/R), GA(L/R), ATL(L/R), CAN(L/R), BU(L/R), AME(L), MB-PED(L/R), MB-CA(L/R) |
| T1 | 39 | BU(L/R), NO, EB, AL(L/R), FLA(R), CAN(L/R), VES(L/R), LOP(R), ATL(L/R), EPA(L/R), PB, LAL(L/R), GA(L/R), GOR(L/R), IPS(L/R), AME(L), ME(R), PLP(R), WED(L), FB, SPS(L/R), AMMC(L/R), IB(L/R), ICL(L), LO(L/R) |
| T2 | 9 | MB-ML(L/R), MB-VL(L/R), MB-CA(R), MB-PED(L/R), CRE(R), LH(R) |
| T3 | 25 | FLA(L), MB-CA(L), AVLP(L/R), SMP(L/R), LH(L), ME(L), SCL(L/R), WED(R), SAD, GNG, AOTU(L/R), PVLP(L/R), ICL(R), PRW, SIP(L/R), SLP(L/R), PLP(L), CRE(L) |

**Supplementary Table 2. Neuropil abbreviations.** Standard three- to four-letter neuropil abbreviations of the Ito et al. reference atlas[3], grouped by anatomical superregion, used throughout the manuscript, figures, and Supplementary Table 1. Suffixes (L), (R), and (L/R) denote the left hemisphere, the right hemisphere, and bilateral pairs, respectively.

| Abbr. | Full name |
|---|---|
| *Optic lobe* | |
| ME | medulla |
| AME | accessory medulla |
| LO | lobula |
| LOP | lobula plate |
| *Central complex* | |
| FB | fan-shaped body |
| EB | ellipsoid body |
| PB | protocerebral bridge |
| NO | noduli |
| *Mushroom body* | |
| MB-CA | calyx |
| MB-PED | peduncle |
| MB-VL | vertical lobe |
| MB-ML | medial lobe |
| *Lateral horn* | |
| LH | lateral horn |
| *Antennal lobe* | |
| AL | antennal lobe |
| *Ventrolateral neuropils* | |
| AVLP | anterior ventrolateral protocerebrum |
| PVLP | posterior ventrolateral protocerebrum |
| PLP | posterior lateral protocerebrum |
| WED | wedge |
| AOTU | anterior optic tubercle |
| *Lateral complex* | |
| BU | bulb |
| LAL | lateral accessory lobe |
| GA | gall |

| Abbr. | Full name |
|---|---|
| *Superior neuropils* | |
| SLP | superior lateral protocerebrum |
| SIP | superior intermediate protocerebrum |
| SMP | superior medial protocerebrum |
| *Inferior neuropils* | |
| CRE | crepine |
| SCL | superior clamp |
| ICL | inferior clamp |
| IB | inferior bridge |
| ATL | antler |
| *Ventromedial neuropils* | |
| VES | vest |
| EPA | epaulette |
| GOR | gorget |
| SPS | superior posterior slope |
| IPS | inferior posterior slope |
| *Periesophageal neuropils* | |
| SAD | saddle |
| FLA | flange |
| CAN | cantle |
| PRW | prow |
| AMMC | antennal mechanosensory and motor center |
| *Gnathal ganglia* | |
| GNG | gnathal ganglia |

**Supplementary Table 3. Cell-type abbreviations.** Cell-type abbreviations used in the Results and Discussion, following the FlyWire adult-brain cell-type annotation[6].

| Abbreviation | Description |
|---|---|
| L1–L5 | lamina monopolar cells |
| C2, C3 | centrifugal cells from medulla to lamina |
| Mi1, Mi4, Mi9 | medulla intrinsic columnar cells |
| Tm1–Tm20 | transmedulla columnar cells |
| TmY3–TmY15 | transmedulla Y-cells |
| T2, T3, T4, T5 | columnar projection neurons (T4 and T5, ON and OFF direction-selective) |
| Pm | proximal-medulla amacrine cells |
| Dm | distal-medulla cells |
| Sm | small-field medulla cells |
| MeMe | medulla-medulla bilateral cells |
| Y1–Y12 | Y-cells |
| CT1 | large amacrine cell |
| Li | lobula intrinsic neurons |
| LMa | lobula multicolumnar amacrines |
| LMt | lobula multicolumnar tangential cells |
| Tlp | translobula-plate cells |
| LPi | lobula-plate intrinsic neurons |
| LC, LPLC | lobula columnar projection neurons |
| ER, ER3a, ER3d, ER5, ER2, ER3m, ER4m | ellipsoid-body ring neurons |
| Delta7 | central-complex Delta neurons |
| KC$\gamma$-m | $\gamma$-main Kenyon cell |
| KC$\alpha\beta$ | $\alpha\beta$ Kenyon cell |
| APL | anterior paired lateral neuron |
| OA-AL2b2 | octopaminergic AL2b2 neuron |
| R7 | chromatic photoreceptor |

Numerical or letter suffixes (e.g., Pm03, Pm08, T5a–d, Tm5c, Tlp4, LMa2, Li10) denote cell-type subtypes as defined by the same annotation[6].

#### Supplementary Note 1: Baseline and control architectures

To isolate the contribution of the connectome scaffold, we compared the connectome-constrained model against two families of generic baselines. Both families shared the same observables (the 73-dimensional neuropil rates), the same train/test split, the same MSE loss (Eq. (7)), and the same Adam[7] schedule, but carried no connectome structure. The two families correspond to the two canonical paradigms for fitting neural-population activity[8,9]: a recurrent generative model that produces trajectories *de novo* from intrinsic noise, and a feedforward next-step predictor that learns the one-step transition operator.

**Gated recurrent unit (GRU).** A standard GRU[10] served as the recurrent generative baseline, an unconstrained recurrent network with no anatomical prior. The hidden state had dimensionality $h = 2{,}048$, and the per-step stochastic drive was a $d_{\text{in}} = 382$-dimensional i.i.d. Gaussian noise vector; these dimensions yield a total parameter count of $\approx 1.5 \times 10^7$, matching the connectome-constrained model within $\pm 5\%$. The GRU was initialized from a learned initial hidden state (analogous to $r_i(0)$ for the connectome-constrained model) and rolled out for the full training duration, producing a 73-dimensional neuropil prediction at each step through a learned linear output projection. Training minimized the same MSE on neuropil rates as the connectome-constrained model. At test time the same learned initial state was used: the GRU was rolled forward over the training segment and then continued autoregressively over the held-out test segment, where predictive metrics were computed.

**Feedforward networks for next-step prediction.** Multilayer perceptrons (MLPs)[11] served as the feedforward baseline, trained under the standard *next-step prediction* paradigm in data-driven dynamics modeling[9]: given the empirical neuropil state $\hat{F}(t) \in \mathbb{R}^{73}$ at time $t$, the network $g_\theta$ predicts the state one imaging step ahead,

$$\begin{aligned} \widetilde{F}(t+\Delta t) &= g_\theta\big(\hat{F}(t)\big), \\ \mathcal{L}_{\text{MLP}} &= \frac{1}{T_{\text{train}}-1}\sum_t \big\|\hat{F}(t+\Delta t) - g_\theta(\hat{F}(t))\big\|_2^2, \end{aligned} \tag{1}$$

with $\hat{F}(t)$ the deconvolved empirical rate of Eq. (2). The total trainable-parameter count of the MLP was matched (within $\pm 5\%$) to that of the connectome-constrained model ($\approx 1.5 \times 10^7$); depth and width were matched between the two MLP variants of Fig. 2k,l so that the only architectural difference was the readout. After fitting, the MLP was evaluated in two modes: a *one-step* mode that scores Eq. (1) on held-out test data, and an *autoregressive rollout* mode that iterates $\widetilde{F}^{(n+1)} = g_\theta(\widetilde{F}^{(n)})$ from an empirical seed state to generate full test-set trajectories for the same distributional and dynamical comparisons used on the connectome-constrained model. Two MLP variants tested the contribution of the connectome readout (Fig. 2k,l): an unconstrained variant whose final layer is a freely learned linear projection onto the 73 neuropils, and a connectome-readout variant whose final layer is replaced by the fixed synapse-density-weighted projection of Eq. (6), leaving only the upstream layers trainable.

GRUs and MLPs are standard architectures; their internal equations are not reproduced here. All baselines used the same random-seed regimen, training-loss-plateau-based stopping, and reporting conventions as the connectome-constrained model.

#### Supplementary Note 2: Avalanche detection and power-law fitting

Cortical resting activity is organized into *neuronal avalanches*, cascades of supra-threshold events whose size and duration follow power-law statistics consistent with dynamics near criticality[12–15]. Throughout, we use "critical" in the operational sense of Sethna and colleagues[16,17]: avalanche sizes and durations are power-law distributed over $\geq 1$ decade, the truncated power law is preferred over lognormal and exponential alternatives by Vuong-corrected likelihood-ratio tests, and the exponents satisfy the crackling-noise scaling relation $(\alpha_S - 1)/(\alpha_D - 1) = 1/(\sigma\nu z)$ within the range reported for near-critical neural recordings. Avalanches are defined on a binary event raster, while the empirical signal is a continuous deconvolved firing-rate proxy and the model output is a continuous rate. Both were mapped to a common event-raster representation by an identical thresholding procedure (below), and the same avalanche definition and statistical tests were applied to the two rasters.

**Event detection.** For both the empirical and the simulated traces, the continuous per-unit activity signal $x_i(t)$ was converted to a binary event raster $b_i(t) \in \{0, 1\}$ following Ref. [14]: each unit's fluctuations were thresholded at three times the standard deviation $\sigma_i$ of its baseline fluctuations[18], set to 1 above threshold and to 0 otherwise,

$$b_i(t) \;=\; \begin{cases} 1, & x_i(t) > 3\sigma_i, \\ 0, & \text{otherwise.} \end{cases} \tag{2}$$

The signal $x_i(t)$ is the deconvolved empirical neuropil rate $\hat{F}_k(t)$ from Eq. (2) for empirical neuropil-level analyses (Fig. 1i; Supplementary Fig. 2a), the simulated neuropil rate $R_k(t)$ from Eq. (6) for simulated neuropil-level analyses (Fig. 1j; Supplementary Fig. 2b), and the simulated single-neuron rate $r_i(t)$ from Eq. (3) for the simulated single-neuron analysis (Fig. 1k; Supplementary Fig. 2c). The bin width was set to $\Delta t_{\text{aval}} = 1/1.2\,\text{s}$, matching the imaging volume rate, for both empirical and simulated rasters. Avalanche analyses were performed exclusively on the held-out test segment for both empirical and simulated traces.

**Avalanche definition.** An avalanche is a contiguous run of population activity. For neuropil-level analyses the per-bin population event signal is the logical OR across the 73 neuropils,

$$A(t) \;=\; \bigvee_i b_i(t), \tag{3}$$

and an avalanche is a maximal interval $[t_{\text{start}}, t_{\text{end}}]$ over which $A(t) = 1$, with $A(t_{\text{start}} - 1) = A(t_{\text{end}} + 1) = 0$. Its duration is

$$D \;=\; t_{\text{end}} - t_{\text{start}} \tag{4}$$

and its size is the total event count across the population during the run,

$$S \;=\; \sum_{t \in [t_{\text{start}}, t_{\text{end}}]} \sum_i b_i(t). \tag{5}$$

For the single-neuron analysis (Fig. 1k; Supplementary Fig. 2c), the same definitions were applied separately to each neuron $i$ by replacing the population OR signal $A(t)$ with the per-neuron event sequence $b_i(t)$: a single-neuron avalanche is a maximal run of $b_i(t) = 1$, with duration $D_i = t_{\text{end}} - t_{\text{start}}$ and size $S_i = \sum_{t \in [t_{\text{start}}, t_{\text{end}}]} b_i(t)$ (the total event count over the active run), so that the single-neuron size and the population size are reported in the same units.

**Power-law fitting and model comparison.** To assess scale-free behavior, we tested whether the empirical and simulated avalanche-duration distributions $P(D)$ and avalanche-size distributions $P(S)$ are consistent with a power law, $P(\cdot) \propto x^{-\alpha}$. The exponent $\alpha$ and the lower cut-off $x_{\text{min}}$ were jointly estimated by maximum-likelihood fitting of a truncated power law[1] in its discrete form (since both $D$ and the population size $S$ are integer-valued in units of $\Delta t_{\text{aval}}$ or event counts); the single-neuron size $S_i$, being a real-valued integral, was fit with the continuous form. Goodness of fit was reported as the Kolmogorov–Smirnov (KS) statistic between the empirical and fitted distributions. To guard against spurious power-law calls, the truncated power law was compared against lognormal and exponential alternatives by likelihood-ratio tests with Vuong's correction for non-nested models (Supplementary Note 3). The same framework was applied to the trained synaptic-weight magnitudes (Fig. 1h), fit with the continuous form; we report the best-fitting lognormal model and its likelihood-ratio comparisons against the power-law and exponential alternatives. All fits and likelihood-ratio tests were performed with the `powerlaw` Python package[2].

#### Supplementary Note 3: Vuong-corrected likelihood-ratio tests for the distributional comparisons of Fig. 1h–k

The distributional comparisons reported in Fig. 1h (absolute synaptic weights against power-law and exponential alternatives) and Fig. 1i–k (avalanche durations against lognormal and exponential alternatives) use the model-selection procedure of Clauset et al.[1] with Vuong's correction for non-nested models[2,19]. For two candidate distributions $\mathcal{L}_1$ and $\mathcal{L}_2$ fit to the same $N$ samples $\{x_n\}$, the log-likelihood-ratio statistic is the per-sample sum

$$R = \sum_{n=1}^{N} \left[ \log \mathcal{L}_1(x_n) - \log \mathcal{L}_2(x_n) \right], \tag{6}$$

and its standardized form

$$z = \frac{R}{\sqrt{N}\,\hat{\sigma}_R} \tag{7}$$

is asymptotically standard normal under the null hypothesis that the two distributions fit the data equally well, where $\hat{\sigma}_R$ is the sample standard deviation of the per-sample log-likelihood differences. The reported two-sided $p$ value is $p = 2\,[1 - \Phi(|z|)]$ with $\Phi$ the standard normal cumulative distribution. The implementation in the `powerlaw` Python package[2] evaluates $\log p$ directly through the asymptotic expansion of $\log \Phi(-|z|)$, so that the test remains well defined when $p$ itself underflows IEEE-754 double precision.

Two consequences shape how the $p$ values reported in Fig. 1h–k should be read. First, $z$ scales as $\sqrt{N}\,\Delta$, with $\Delta$ the typical per-sample log-likelihood gap between the two candidate distributions. At $N \approx 1.5 \times 10^7$ synapses (Fig. 1h), a per-sample gap of order $10^{-3}$ in natural-log units already drives $|z|$ into the regime where $\Phi(-|z|)$ falls below the double-precision floor of $\sim 10^{-308}$. The package-internal log-space values are then on the order of $-10^4$ for the lognormal-versus-exponential comparison and $-10^7$ for the lognormal-versus-power-law comparison, both of which exceed the dynamic range of standard scientific notation and are not interpretable as conventional $p$ values. We therefore report Fig. 1h as $p < 10^{-300}$ for both alternatives rather than quoting the package's literal log-space output. The same bound applies to the exponential-alternative comparisons referenced in Fig. 1i–k, which are dominated by the same sample-size effect and are systematically smaller than the lognormal-alternative $p$ values displayed in those panels.

Second, this $\sqrt{N}$ scaling decouples statistical significance from goodness of fit at the sample sizes used here. A $p$ value satisfying $p < 10^{-7}$ at $N \sim 10^7$ is not evidence that the preferred distribution matches the data accurately; it indicates only that the alternative is worse by an amount the test can resolve. The practical fit quality should be read off the Kolmogorov–Smirnov statistic ($KS$) and the explicit exponent reported in each panel of Fig. 1h–k, not off $p$. For this reason, and for the paired $t$-tests across $n = 5$ trained models reported in Figs. 4i,j,k,l and 5p–s, the manuscript reports numerical $p$ values without categorical significance asterisks; n.s. denotes $p \geq 0.05$.

#### Supplementary Note 4: Identifiability of the fitted effective synaptic weights

The connectome-constrained model learns roughly $1.5 \times 10^7$ synaptic-weight magnitudes $\{|w_{ij}|\}$, together with 138,639 per-neuron time constants, 138,639 per-neuron initial rates, and two global drive parameters (Eq. (3)), against the 73-dimensional neuropil time series of a single recording half (Eq. (7)). By parameter count alone the inverse problem is underdetermined, so the fitted weights could be any point in a large set of equally good solutions rather than a meaningful estimate. Yet every mechanistic result, from cluster-level necessity to the inhibitory hubs and their cell-type silencing, depends on the weights carrying information. We therefore set out what the fit does and does not identify, using the reproducibility controls of Fig. 2m–p and Supplementary Fig. 4.

The parameter count overstates the freedom in the fit, because the connectome fixes three things the optimizer never adjusts. Support: a weight exists only where two neurons are synaptically connected, about $1.5 \times 10^7$ of the $\sim 1.9 \times 10^{10}$ ordered pairs among 138,639 neurons. Anatomy therefore pins more than 99.9% of the weight matrix to zero (Connectome data and neuron–synapse assignment). Sign: each neuron's polarity is frozen from the FlyWire neurotransmitter prediction[6], so only the magnitude $|w_{ij}|$ is learned, and all of a neuron's outgoing weights share one sign (Eq. (3)). Readout: the projection from 138,639 single-neuron rates onto the 73 observed neuropil channels is the fixed synapse-density average of Eq. (6). It has no free parameters, so the map from latent to observed variables is set by anatomy, not by the fit (Connectome-derived neuropil readout). These constraints do not make the problem formally well-posed, but they tie the high-dimensional weights to the low-dimensional target through a fixed structure. That structure lets a 73-channel recording shape $10^7$ magnitudes toward a repeatable solution.

Three controls test whether that solution is repeatable. Across initialization families, a uniform interval and a truncated-normal prior centered at zero converge to statistically indistinguishable weight distributions (Fig. 2o,p). Their neuropil-aggregated excitatory and inhibitory weights agree to within ±5% (Supplementary Fig. 4b). Across random seeds at a fixed prior, agreement is again within ±5% (Supplementary Fig. 4c). Across the gradient estimator, replacing backpropagation through time with the online propagator of BrainTrace[20] yields fits of comparable quality (Training data, loss function, and optimization), so the solution does not depend on how gradients are computed. This agreement is not inherited from a shared starting point. Training moves the weights far from every initialization (Fig. 2m,n; Supplementary Fig. 4a) before they converge on the same aggregates.

Reproducibility matters at the level where the claims are made. No result rests on the exact magnitude of a single synapse. The claims are distributional (the lognormal weight law, Fig. 1h), cluster-level (the necessity of the S1 core), and cell-type-level (the inhibitory hubs), and the quantities behind them inherit the reproducibility shown above. The directed effectome, computed from the learned weights, returns the same source (S1, S2, S3) and target (T1, T2, T3) partitions across the $n = 5$ independently trained models. Its adjusted Rand index against the averaged clustering is well above chance (Supplementary Fig. 6; Neuropil effectome construction and clustering). The aggregates the paper interprets are therefore identified, even though the individual weights inside them are too numerous for the data to constrain one by one.

Three boundaries follow. First, individual per-synapse magnitudes are not separately identified. Reproducibility holds at the neuropil-aggregate, distributional, and cluster levels, and the paper makes no claim about any single synapse's exact value. Second, the absolute scale is conventional, fixed by rescaling the deconvolved target to a mean rate of 1 Hz (Calcium-to-firing-rate deconvolution), so only relative weights are interpretable. Third, all main-text fits come from one fly, so the cross-model agreement measures fitting variability, not biological variability. Identifiability across animals is untested, a prediction for multi-animal connectome-plus-recording data (Discussion).

#### Supplementary References


1. Clauset, A., Shalizi, C. R. & Newman, M. E. J. Power-law distributions in empirical data. *SIAM Review* **51**, 661–703, DOI: 10.1137/070710111 (2009).
2. Alstott, J., Bullmore, E. & Plenz, D. powerlaw: a Python package for analysis of heavy-tailed distributions. *PLoS ONE* **9**, e85777, DOI: 10.1371/journal.pone.0085777 (2014).
3. Ito, K. *et al.* A systematic nomenclature for the insect brain. *Neuron* **81**, 755–765, DOI: 10.1016/j.neuron.2013.12.017 (2014).
4. Fox, M. D. & Raichle, M. E. Spontaneous fluctuations in brain activity observed with functional magnetic resonance imaging. *Nature reviews neuroscience* **8**, 700–711, DOI: 10.1038/nrn2201 (2007).
5. Smith, S. M. *et al.* Correspondence of the brain's functional architecture during activation and rest. *Proceedings national academy sciences* **106**, 13040–13045, DOI: 10.1073/pnas.0905267106 (2009).
6. Schlegel, P. *et al.* Whole-brain annotation and multi-connectome cell typing of Drosophila. *Nature* **634**, 139–152, DOI: 10.1038/s41586-024-07686-5 (2024).
7. Kingma, D. P. & Ba, J. Adam: a method for stochastic optimization. In *Proceedings of the 3rd International Conference on Learning Representations (ICLR)* (2015).
8. Sussillo, D. & Abbott, L. F. Generating coherent patterns of activity from chaotic neural networks. *Neuron* **63**, 544–557, DOI: 10.1016/j.neuron.2009.07.018 (2009).
9. Pandarinath, C. *et al.* Inferring single-trial neural population dynamics using sequential auto-encoders. *Nature Methods* **15**, 805–815, DOI: 10.1038/s41592-018-0109-9 (2018).
10. Cho, K. *et al.* Learning phrase representations using RNN encoder–decoder for statistical machine translation. In *Proceedings of the 2014 Conference on Empirical Methods in Natural Language Processing (EMNLP)*, 1724–1734, DOI: 10.3115/v1/D14-1179 (2014).
11. Goodfellow, I., Bengio, Y. & Courville, A. *Deep Learning* (MIT Press, Cambridge, MA, 2016).
12. Beggs, J. M. & Plenz, D. Neuronal avalanches in neocortical circuits. *Journal neuroscience* **23**, 11167–11177, DOI: 10.1523/JNEUROSCI.23-35-11167.2003 (2003).
13. Shriki, O. *et al.* Neuronal avalanches in the resting MEG of the human brain. *Journal Neuroscience* **33**, 7079–7090, DOI: 10.1523/JNEUROSCI.4286-12.2013 (2013).
14. Ponce-Alvarez, A., Jouary, A., Privat, M., Deco, G. & Sumbre, G. Whole-brain neuronal activity displays crackling noise dynamics. *Neuron* **100**, 1446–1459.e6, DOI: 10.1016/j.neuron.2018.10.045 (2018).
15. Nanda, A. *et al.* Time-resolved correlation of distributed brain activity tracks EI balance and accounts for diverse scale-free phenomena. *Cell reports* **42**, DOI: 10.1016/j.celrep.2023.112254 (2023).
16. Sethna, J. P., Dahmen, K. A. & Myers, C. R. Crackling noise. *Nature* **410**, 242–250, DOI: 10.1038/35065675 (2001).
17. Friedman, N. *et al.* Universal critical dynamics in high resolution neuronal avalanche data. *Physical Review Letters* **108**, 208102, DOI: 10.1103/PhysRevLett.108.208102 (2012).

18. Romano, S. A. *et al.* An integrated calcium imaging processing toolbox for the analysis of neuronal population dynamics. *PLoS Computational Biology* **13**, e1005526, DOI: 10.1371/journal.pcbi.1005526 (2017).
19. Vuong, Q. H. Likelihood ratio tests for model selection and non-nested hypotheses. *Econometrica* **57**, 307–333, DOI: 10.2307/1912557 (1989).
20. Wang, C. *et al.* Model-agnostic linear-memory online learning in spiking neural networks. *Nature Communications* DOI: 10.1038/s41467-026-68453-w (2026).
